# Thermal biology diversity of bee pollinators: taxonomic, phylogenetic and plant community-level correlates

**DOI:** 10.1101/2023.11.27.568883

**Authors:** Carlos M. Herrera

## Abstract

Community-wide assembly of plant-pollinator systems depends on an intricate combination of biotic and abiotic factors, including heterogeneity among pollinators in thermal biology and responses to abiotic factors. Studies on the thermal biology of pollinators have mostly considered only one or a few species of plants or pollinators at a time, and the possible driving role of the diversity in thermal biology of pollinator asemblages at the plant community level remains largely unexplored. More specifically, it is unknown whether diversity in the thermal biology of bees, a major pollinator group worldwide, contributes to the assembly and maintenance of diverse bee communities, broadens the spectrum of possibilities available to bee-pollinated plants, facilitate interspecific partitioning of ecological gradients across habitats, seasons and time of day, and/or enhance plant pollination success through complementarity effects. The objectives of this study were to assess the diversity in thermal biology of the bee assemblage that pollinates plants in a Mediterranean montane area, evaluate its taxonomic and phylogenetic underpinnings, and elucidate whether there existed seasonal, daily, between-habitat or floral visitation correlates of bee thermal biology which could contribute to partition ecological gradients among plant and bee species. Thermal biology parameters were obtained in the laboratory (*K*, intrinsic warming constant) and the field (thoracic and ambient temperature at foraging site, *T*_th_ and *T*_air_) on individual bees of a diverse sample (*N* = 204 bee species) comprising most bee pollinators of the regional plant community. Species-specific thermal biology parameters were combined with quantitative field data on bee pollinators and flower visitation for the regional community of entomophilous plants (*N* = 292 plant species). Results revealed that the regional bee assemblage harbored considerable diversity in thermal biology features, that such diversity was mostly taxonomically, phylogenetically and body-size structured, and that the broad interspecific heterogeneity in thermal biology represented in the bee community as a whole eventually translated into daily, seasonal, among-habitat and flower visitation patterns at the plant community level. This lends support to the hypothesis that broad diversity in thermal biology of bees can act enhancing opportunities for bee coexistence, spatio-temporal partitioning of floral resources, and plant pollination success.

## INTRODUCTION

From the viewpoint of animal consumers, floral resources are typically arranged as discrete, ephemeral, sparse, individually little-rewarding items. From the plants’ perspective, these features contribute to enhance the movement of pollen between individuals and reduce self-pollination (Dreisig 1985, Harder and Barrett 1995, Biernaskie et al. 2002, Ferdy and Smithson 2002). Pollination occurs as a merely incidental consequence of the “sloppy harvest” (Janzen 1983) of floral resources by foraging animals, which have to spend considerable time and energy in the process to meet their own food and energy requirements and, in the extreme case exemplified by bees, also for provisioning their offspring (Heinrich 1975, Michener 2000, McCallum et al. 2013). An evolutionary tension or antagonism (”evolutionary game”, Pyke 2016) thus arises consisting of conflicting selective pressures: on the plants, to evolve features that augment the amount of (costly) movement needed by pollinators to fulfill their energetic requirements (e.g., by reducing the mean and/or increasing the variance among flowers in amount or quality of food reward; Keasar et al. 2008, Herrera 2009, Pyke et al. 2020, Maldonado et al. 2023); and on the pollinators, to diminish the energetic costs associated with the movement between, and handling of, food units (e.g., by optimizing departure rules from flowers in function of reward availability and distribution; Pyke 1979, 2016, Kadmon and Shmida 1992). Consideration of these aspects motivated the formulation of an energetics-oriented approach to pollination ecology that emphasized, among other, the crucial roles of pollinators’ energy requirements, energy expenditure, and thermal biology features as major drivers of the assembly of plant-pollinator interactions in time and space (Heinrich 1975, 1981, Heinrich and Raven 1972, Willmer 1983, Abrol 2012, McCallum et al. 2013, Pyke 2016, Balfour et al. 2021, Herrera et al. 2023; among many others).

Most previous studies on the energetics of pollination or the thermal biology of pollinators have considered only one or a few species of plants or pollinators at a time (Herrera 1995, 1997, Stone et al. 1999, Orueta 2002, Nieh et al. 2006, Seymour et al. 2009, Corbet and Huang 2016, Balfour et al. 2021, Leclerc et al. 2022; see Abrol 2012, McCallum et al. 2013 for reviews). Few investigations have attempted so far to relate the thermal biology of broad pollinator groups with the pollination features of plant communities (Warren et al. 1988, Shmida and Dukas 1990, Herrera 1997, Osorio-Canadas et al. 2016, Dellinger et al. 2021), and even fewer have actually undertaken empirical work on the thermal biology of pollinator asemblages at the community level (Herrera et al. 2023). In a Mediterranean montane plant community predominantly pollinated by bees (Herrera 2020), Herrera et al. (2023) recently found that the seasonal activity of *Andrena* bees and their predominant role in the pollination of early-blooming plants can be parsimoniously explained by features of their thermal biology: null to weak endothermy, ability to forage at low body temperature, low upper tolerable limit of body temperature, weak thermoregulatory capacity, and high warming constant enhancing ectothermic warming. These results contrasted with those of previous work on the thermal biology of bees from other families, which had mostly documented strong endothermy and high body temperatures (May and Casey 1983, Stone and Willmer 1989, Stone 1993a,b, Heinrich 1993, Roberts et al. 1998, Oyen et al. 2016). These seemingly conflicting results, along with those of Stone (1994) and Bishop and Armbruster (1999), actually suggest a strong phylogenetic component in bee thermal biology, which in turn highlights the need of evaluating the diversity in thermal biology represented in taxonomically diverse bee pollinator assemblages and its potential ecological consequences in relation to the pollination of entire plant communities. Bees are the dominant pollinators in many plant communities worldwide (Ballantyne et al. 2017, Herrera 2020, Simpson et al. 2022). To the extent that relevant aspects of the bees’ thermal biology are interspecifically variable and determine their daily or seasonal periods of activity, habitat use, flower preferences, or flower visitation rates via response diversity effects (Fründ et al. 2013a), community-wide diversity of bee thermal biology would facilitate interspecific partitioning of ecological gradients occurring over habitat types, seasons and/or time of day. Through this little explored mechanism, thermal diversity in itself could contribute to the assembly and maintenance of diverse bee communities, broaden the spectrum of possibilities available to bee-pollinated plants, and enhance overall plant pollination success by promoting complementarity effects (Willmer and Corbet 1981, Albrecht et al. 2012, Fründ et al. 2013b, Simpson et al. 2022).

The main objectives of this study were (1) to evaluate the diversity in thermal biology features of the very species-rich bee assemblage that pollinates the diverse community of entomophilous plants inhabiting a well-preserved Mediterranean montane area; (2) to assess the taxonomic and phylogenetic underpinnings of such diversity; and (3) to elucidate whether heterogeneity in the thermal biology of bee assemblages had discernible seasonal, daily, between-habitat, and/or floral visitation correlates which would support the hypothesis posited above (see also Willmer and Corbet 1981) that diversity in thermal biology can ultimately contribute to partition spatio-temporal ecological gradients among bee and plant species. To address the first objective, thermal biology parameters were experimentally obtained in the laboratory (intrinsic warming constant) or measured in the field (paired thoracic and ambient temperatures) on a large sample of individual bees. To properly evaluate the taxonomic and phylogenetic basis of the diversity in thermal biology of the regional bee assemblage (Objective 2), efforts were made to sample as many and taxonomically diverse bee species as possible. Thermal biology data were gathered for most bee pollinators of the regional plant community (*N* = 204 bee species), which also accounted for a substantial fraction of the regional bee fauna. To assess whether diversity in thermal biology of bees contributed to create spatial, temporal or flower visitation patterns (Objective 3), species-specific thermal biology parameters were merged with detailed quantitative data on bee pollinator composition and flower visitation for virtually every entomophilous plant species occurring in the study region (*N* = 292 plant species). Results revealed that bee communities harbored considerable diversity in thermal biology features, that such diversity was mostly taxonomically, phylogenetically, and body-size structured, and that the broad interspecific heterogeneity in thermal biology represented in the bee community eventually translated into daily, seasonal, among-habitat and flower visitation patterns by bees at the plant community level.

## MATERIALS AND METHODS

### Study area and sampling periods

Thermal biology parameters were measured in the field or assessed in the laboratory on wild bees sampled in the Sierra de Cazorla (Jaén Province, southeastern Spain) between July 1990– October 2023. Standardized pollinator censuses that provided the raw data used here to characterize the composition of bee pollinator assemblages with reference to their thermal biology were conducted in the same area between July 1997–August 2023. A map of sampling sites is shown in Herrera (2021), and a series of representative photographs depicting major habitat types in a subset of sampling locations can be found in Herrera (2019: Appendix S1). The Sierra de Cazorla area is characterized by well-preserved montane Mediterranean habitats and outstanding biological diversity (Médail and Diadema 2009, Gómez Mercado 2011). In particular, the wild bee fauna of the area, with 392 species recorded to date, is strikingly diverse in comparison with other Mediterranean regions (see Ortiz-Sánchez et al. 2023 for comparisons). For some bee species which were rare or difficult to sample in the study area, field measurements (21 % of total field measurements) were taken in February-April 2022 at two locations in Sierra Morena (Córdoba province) and the lowlands of the Guadalquivir River valley (Sevilla province), 160 km and 300 km away, respectively, from the Cazorla main study area. Analytical results obtained with and without these data were essentially identical, which warranted its inclusion for enhancing statistical power and taxonomic/phylogenetic completeness of the sample.

### Bee thermal biology

#### Field measurements

Free-ranging bees were hand netted in fine weather while foraging at flowers of different plants. Bees were sampled over as broad ranges as possible of time of year, habitat type and ambient temperature. For each bee, thoracic temperature (*T*_th_) was measured within 10–20 s of netting to the nearest 0.1°C using a fast-response (time constant 0.025 s), 0.33 mm-diameter needle microprobe with sharpened tip (Type MT-29/1; Physitemp Instruments, Clifton, New Jersey, USA). Readings were obtained by inserting the probe 1 mm into the dorsal midline of the bee’s thorax while it was restrained in the net. Air temperature (*T*_air_) 2-5 cm away from the flower where the bee had been caught was measured within 2 min of the bee’s capture using a digital thermometer and a fast-response 0.22 mm-diameter thermocouple. A total of 1,936 pairs of temperature measurements for 95 bee species in 21 genera were taken in the field on 84 different dates (see Appendix S1: Table S1 for bee species and sample sizes). Preliminary assays showed that insertion of the needle microprobe into the bees’ thorax produced unreliable *T*_th_ measurements and undesirable bee mortality when applied to individuals with body mass under ∼20 mg. For these reasons, thermal data were not gathered for these species in the field, which resulted in very small bees (typically *Hylaeus*, *Halictus* subgen. *Seladonia*, *Lasioglossum* subgen. *Dialictus*, *Andrena* subgen. *Micrandrena*) being underrepresented in the set of *T*_th_ and *T*_air_ data relative to their frequency in the bee fauna.

#### Laboratory experiments

The intrinsic warming constant (*K*, see *Data analysis* below for definition) of individual bees belonging to different species was assessed experimentally on specimens netted in the field while foraging at flowers. They were placed into sealed microcentrifuge tubes after capture, kept in the dark in a portable refrigerator, and quickly brought to the laboratory. Within 6 h of capture, bees were weighed, killed using ethyl acetate, and had the junction of a fast-response 0.22mm Type T thermocouple implanted in the thorax to a depth of 1 mm. The dead bee was cooled in a refrigerator until its thoracic temperature was ∼4-6°C, then suspended in the air 2-3 cm above a piece of styrofoam in a small room with still air (room temperature 16-22°C) and allowed to passively warm up until stabilization of thoracic temperature (*T*_th_). During this period, *T*_th_ and air temperature 3 cm away from the bee were recorded every 2 s using a Omega HH520 data logger/thermometer. I applied this experimental protocol to 645 individuals from 169 bee species, 30 genera and five families collected on 99 different dates. The temporal series of thermal data thus obtained were used to obtain quantitative estimates of their intrinsic warming constant (see *Data analyses* below). The full range of body sizes occurring in the regional bee community, including many small species not amenable to *T*_th_ measurements, was included in the experiments for estimating *K* values (Appendix S1, Table S2).

### Pollinator censuses

#### Plant community sampling

Pollinator composition was quantitatively assessed for 292 plant species in 187 genera from 54 families, which are a superset of the 191 and 221 species studied by Herrera (2020 and 2021, respectively), and include virtually all entomophilous plants occurring in the Sierra de Cazorla area. Mean pollinator sampling date for each plant species roughly matched its peak flowering date, hence the seasonal distribution of sampling times closely matched the pattern of flowering times in the region (see Herrera 2019, 2020, for additional details).

Obtaining quantitative pollinator data for a large species sample virtually encompassing the whole plant community required spanning field work over many years, since pre-established replication rules (see *Census methods* below) limited the number of species that could be sampled per year. Pollinator sampling was conducted on a single site in the vast majority of the species considered here (*N* = 274), while 18 species were sampled on two or more sites. Pollinator composition of some plant species can vary among sites or years, but such intraspecific variation is quantitatively minor in comparison to the broad interspecific variation occurring in the large species sample considered here, as previously shown by Herrera (2020: Table 2 and Fig. 2) by applying formal variance partitions. Pollinator data for the same plant species obtained in more than one site or year were thus combined into a single sample for the analyses. The quality of pollinator composition data as assessed with Good-Turing’s “sample coverage” parameter, which estimates the proportion of the total population of pollinators that is represented by the species occurring in the sample (Good 1953, Hsieh et al. 2016), was very high (>85%) for the vast majority of plant species (see Appendix S1: Table S1 in Herrera 2021).

To asess possible habitat correlates of the thermal biology of bee pollinator assemblages, each plant species with pollinator census data was assigned to one of the following nine habitat types (total species per habitat in parentheses): vertical rock cliffs (20 species); local disturbances caused by humans, large mammals or natural abiotic processes (28); sandy or rocky dolomitic outcrops (44); dwarf mountain scrub dominated by cushion plants (38); forest edges and large clearings (39); deciduous, mixed or coniferous forest interior (26); patches of grasslands and meadows on deep soils in relatively flat terrain (51); tall, dense Mediterranean sclerophyllous forest and scrub (22); banks of permanent streams or flooded/damp areas around springs (24). Species that occurred in more than one of these habitat types were assigned to the habitat where the pollinator censuses were conducted.

#### Census methods

Data on bee visitation to flowers were obtained by applying the same standardized sampling protocol for all plant species studied (Herrera 2019, 2020). The sampling unit for pollinator assessment was the “pollinator census”, consisting of a 3-min watch of a flowering patch whose total number of open flowers was also counted. Bees visiting some flower in the focal patch during the 3-min period were identified (see *Bee identification* below), and total number of flowers probed by each individual was recorded. Areal extent and number of open flowers in monitored patches were adjusted for each plant species according to flower size and density so that I could confidently monitor all pollinator activity in the patches from a distance of 1.5-2 m. Assessing the number of elemental florets visited by pollinators was impractical in species with tiny flowers densely packed into compact inflorescences (e.g., Apiaceae, Asteraceae; *N* = 59 species). In these cases the number of inflorescences available per patch and visited per census by individual bees were counted rather than individual flowers, and flower visitation rates (see *Data analysis* below) thus actually referred to inflorescences. I will collectively refer to visitation to both single flowers and inflorescences as “flower visitation” for simplicity.

Census replication rules for each plant species-site-year combination involved a minimum of 60 censuses spread over three non-consecutive dates and conducted on ≥20 widely spaced flowering patches with roughly similar flower numbers. On each date, censuses should be distributed from 0.5-2.5 h past sunrise (depending on season; censuses started earlier in summer) through one hour past noon, the different patches being watched in random order. Flowers of about half of the species studied are not available to pollinators in the afternoon, as their corollas wither, close or fall shortly after noon, and earlier studies in the area have also shown that insect pollinator activity declines considerably in the afternoon (Herrera 1990, 1995, C. M. Herrera *unpublished observations*). Data used in this study on the composition and abundance of the bee pollinator assemblage were based on a total of 36,457 pollinator censuses carried out on 763 different dates and accounting for a total watching effort of 8,092,356 flower·min.

#### Bee identification

Bees recorded during pollinator censuses were identified by combining specimen collection with systematic photographic recording. Close-up photographs of bees visiting flowers were taken routinely during censuses using a DSLR digital camera and 105 mm macro lens. These photographs were used for identification, keeping photographic vouchers, and ascertaining the pollinating status of different taxa. Bee specimens were also collected regularly for identification or confirmation by specialists. Bee taxonomists that contributed identifications are listed in *Acknowledgments*. Out of a total of 18,622 individual bees recorded in censuses, 95.4% were confidently identified to species, 1.2% were assigned to cryptic species pairs of congeneric species, and 3.4% could be identified only to genus. All bee taxa recorded in censuses were considered here as pollinators, as they contacted anthers or stigmas, and/or visibly carried pollen grains on body surfaces.

### Data analysis

#### Thermal biology diversity

All statistical analyses reported in this paper were carried out using the R environment (R Core Team 2023). Data from the laboratory runs on dead bees were used to estimate their intrinsic thermal constant *K*, which described heat transfer rate and were obtained by fitting the passive warming curve of the dead bee to the equation *dT*/*dt* = *K* (*T*-*T*_air_) (Newton’s law of cooling; Bakken 1976, Willmer and Unwin 1981, Casey 1988), in which *T* is the temperature of the object, *t* is time, *T*_air_ is ambient temperature, and *K* is the cooling/warming constant (”warming constant” hereafter). The nonlinear least-squares method implemented in the function nls of the stats package was used to estimate *K* from the empirical experimental data. *K* values were used to estimate the intrinsic heat transfer rates of the species tested. Independent replicates obtained for different individuals of the same species were averaged to obtain a single *K* value per species. See Appendix S1: Table S2 for bee species, sample sizes, and species means.

Relationships between the thermal parameters (*K*, *T*_th_, *T*_exc_), on one side, and body mass and taxonomic affiliation, on the other, were tested by fitting linear models with species means as response variables and body mass (log transformed), bee family, and bee genus nested within family, as predictors. As strong taxonomic effects on all thermal parameters were found, the relationship between thermal biology diversity and bee phylogeny deserved detailed investigation. By pruning the large, albeit partial (∼23% of world bees) bee supermatrix phylogeny of Henríquez-Piskulich et al. (2024), I constructed two phylogenetic trees comprising, respectively, 125 of the species in my sample with *K* data (74% of total) and 71 of the species in my sample with *T*_th_ and *T*_exc_ data (75% of total). Tests of phylogenetic signal in thermal biology parameters (”tendency for related species to resemble each other more than they resemble species drawn at random from the tree”, Blomberg and Garland 2002) were conducted using Pagel’s λ, which performs well to discriminate between random and Brownian motion patterns of trait distribution; is robust to variations in the number of species in the phylogeny and to incompletely resolved phylogenies or suboptimal branch-length information; and provides a reliable effect size measure (Münkemüller et al. 2012, Molina-Venegas and Rodríguez 2017). To identify relevant “local hotspots” of phylogenetic autocorrelation contributing disproportionately to overall phylogenetic signal in thermal biology parameters, local Moran’s *I* (*I*_i_) was computed for each tip of the phylogenetic tree, which provided an indication of the extent of significant phylogenetic clustering of similar values around that tip (Anselin 1995). Computations were performed using the package phylosignal (Keck et al. 2016), and statistical significance of Pagel’s λ and *I*_i_ was tested by randomization.

#### Ecological patterns

Daily, yearly, and among-habitat variation in the thermal biology profiles of bee pollinators at the bee and plant community levels, as well as the relationships between flower visitation rate, flower type, and thermal biology, were examined by merging relevant plant and bee information at species level into the dataset containing the raw pollinator census data at individual bee level. For the analyses of daily variation, time of sunrise was computed for each pollinator census date using the getSunlightTimes function in the suncalc package (Thieurmel and Elmarhraoui 2022). Each plant species was assigned to one of the nine habitat types (see *Plant community sampling* above) and two perianth classes, the latter for assessing possible relationships between a plant species’ flower type and the thermal biology of its bee pollinators. The two perianth classes recognized corresponded to open, more or less bowl-shaped, non-restrictive perianths (”open perianth” hereafter, *N* = 117 species), and closed, tubular, sympetalous or otherwise restrictive perianths (”restrictive perianth” hereafter, *N* = 175 species). Flower visitation rate was the number of flowers visited during the 3-min census period by each individual bee.

Each bee recorded in censuses was characterized by the species’ means for body mass, warming constant (*K*), thoracic temperature (*T*_th_), and thermal excess (*T*_exc_ = *T*_th_ – *T*_air_) whenever these data were available (Appendix S1, Tables S1 and S2). This procedure was motivated by the fact that intraspecific variance in these parameters (i.e., due to variation among conspecific individuals) was comparatively minor relative to the variance due to differences among families, genera and species combined (range = 73-88%; Appendix S1, Table S3). In total, 9,012 individual bee records from the censuses could be associated with species-specific *T*_th_ and *T*_exc_ values, and 14,921 individuals with species-specific *K* values, out of a total of 18,622 bees recorded in all censuses combined. The lower number of individuals with *T*_th_ and *T*_exc_ data was mainly due to the difficulties involved in taking these measurements on small bees noted above (see *Bee thermal biology, Field measurements*). Body mass differed considerably among bee species (range = 5.1–543.5 mg, interquartile range = 21.4–84.4 mg, *N* = 204 species with body mass data; Appendix S1: Table S2) and the possible confounding effect of this variable was accounted for in the analyses by including it in every model testing for ecological correlates of thermal parameters.

All simple linear models reported in this paper were conducted using the function lm in the stats package, while linear mixed-effect models were fit with the lmer function in package lme4 (Bates et al. 2015). Model fit adequacy was in every instance checked using the check_model function in the performance package (Lüdecke et al. 2021), and transformations were applied to the data whenever they improved model adequacy. The statistical significance of fixed effects and covariates was assessed with the Anova function in the car package (Fox and Weisberg 2019). The function ggpredict from the ggeffects package (Lüdecke 2018) was used to compute marginal effects of thermal parameters on flower visitation rates (i.e., after controlling for the combined effects of the rest of predictors and covariates in the model). Modality of distributions was tested using Hartigans’ dip modality test (Hartigan and Hartigan 1985) as implemented in function dip.test of the diptest package (Maechler 2021). Nonparametric regression methods based on the generalized additive model smoother in function gam of the mgcv package (Wood 2017) were used to examine daily and annual variation in the thermal profile of bee pollinator assemblages, which were expected to depart substantially from simple parametric relationships. Nonparametric regressions, in contrast to ordinary parametric linear or quadratic regressions, do not make *a priori* assumptions on the shape on the statistical relationship linking predictor and response variables (Wood 2017).

## RESULTS

### Bee thermal biology: warming constant

There was extensive interspecific variation in warming constant (*K*), which ranged between 0.00437– 0.07176 s^-1^ (interquartile range = 0.01100–0.02309 s^-1^), or a 16-fold variation. Such broad variation in *K* was statistically accounted for by the combined effects of interspecific variation in body mass and intrinsic features of bee families and genera (Table 1; see Appendix S1: Figure S1 for a plot depicting the inverse relationship betwen *K* and body mass across bee species, all families and genera combined, and Appendix S1: Figure S2 for separate plots by family). Despite broad overlap between families (Figure 1A), interfamilial heterogeneity was statistically significant (Table 1). On average (± SE; this notation will be used hereafter when reporting means), the warming constant declined in the direction Halictidae (0.0373 ± 0.00301, *N* = 28 species), Colletidae (0.0300 ± 0.00658, *N* = 8), Andrenidae (0.0243± 0.00178, *N* = 48), Megachilidae (0.0173 ± 0.00151, *N* = 37) and Apidae (0.0168 ± 0.00188, *N* = 49). The mean warming constant of the family with the fastest heat transfer rate (Halictidae) thus more than doubled that of the family with the slowest transfer rate (Apidae). Differences in *K* among families persisted after accounting for body mass (Table 1, Appendix S1: Figure S2). For a given body mass, Halictidae and Andrenidae tended to have the highest heat transfer rates, and Apidae and Megachilidae the lowest ones. The significant differences in *K* between genera of the same family (after statistically accounting for body mass, Table 1) are well illustrated in Figure 1B by the very genus-rich families Apidae and Megachilidae. In the former, for instance, species of *Ceratina*, *Epeolus* and *Nomada* had consistently higher warming constant than species of *Eucera*, *Amegilla*, *Anthophora*, *Bombus* or *Xylocopa* (Figure 1B). Heterogeneity among bee families, and among genera within families, in thermal constant (*K*) mostly reflected the strong phylogenetic signal underlying the data, as denoted by the large, highly significant Pagel’s λ (Table 2), and the strongly clustered distribution of local Moran’s *I* across the phylogenetic tree of bee species in the sample (Figure 2A).

**TABLE 1.** Summary of results of linear models testing for the effects of mean body mass (mg, log-transformed), bee family, and bee genus (nested within family) on the species means for warming constant (*K*, s^-1^, log-transformed), thoracic temperature (*T*_th_, °C), and thermal excess (*T*_exc_ = *T*_th_ – *T*_air_, °C).

| Response variable | Model $R^2$ | Predictor | Parameter estimate ± SE | Degrees of freedom | $F$ | $P$-value |
| --- | --- | --- | --- | --- | --- | --- |
| $\text{Log}_{10}(K)$ | 0.985 | $\text{log}_{10}(\text{body mass})$ | $-0.63 \pm 0.01$ | 1, 138 | 3427.6 | $<2.2E-16$ |
|  |  | Family |  | 4, 138 | 12.4 | 1.2E-08 |
|  |  | Genus within Family |  | 25, 138 | 3.3 | 4.6E-06 |
| $T_{th}$ | 0.810 | $\text{log}_{10}(\text{body mass})$ | $3.01 \pm 1.18$ | 1,72 | 6.6 | 0.013 |
| | | Family | | 4,72 | 35.6 | $<2.2E-16$ |
|  |  | Genus within Family |  | 16,72 | 4.4 | 5.44E-06 |
| $T_{exc}$ | 0.732 | $\text{log}_{10}(\text{body mass})$ | $6.63 \pm 1.37$ | 1,72 | 23.5 | 6.9E-06 |
|  |  | Family |  | 4,72 | 10.8 | 6.8E-07 |
|  |  | Genus within Family |  | 16,72 | 3.3 | 0.00031 |

**Figure 1.**
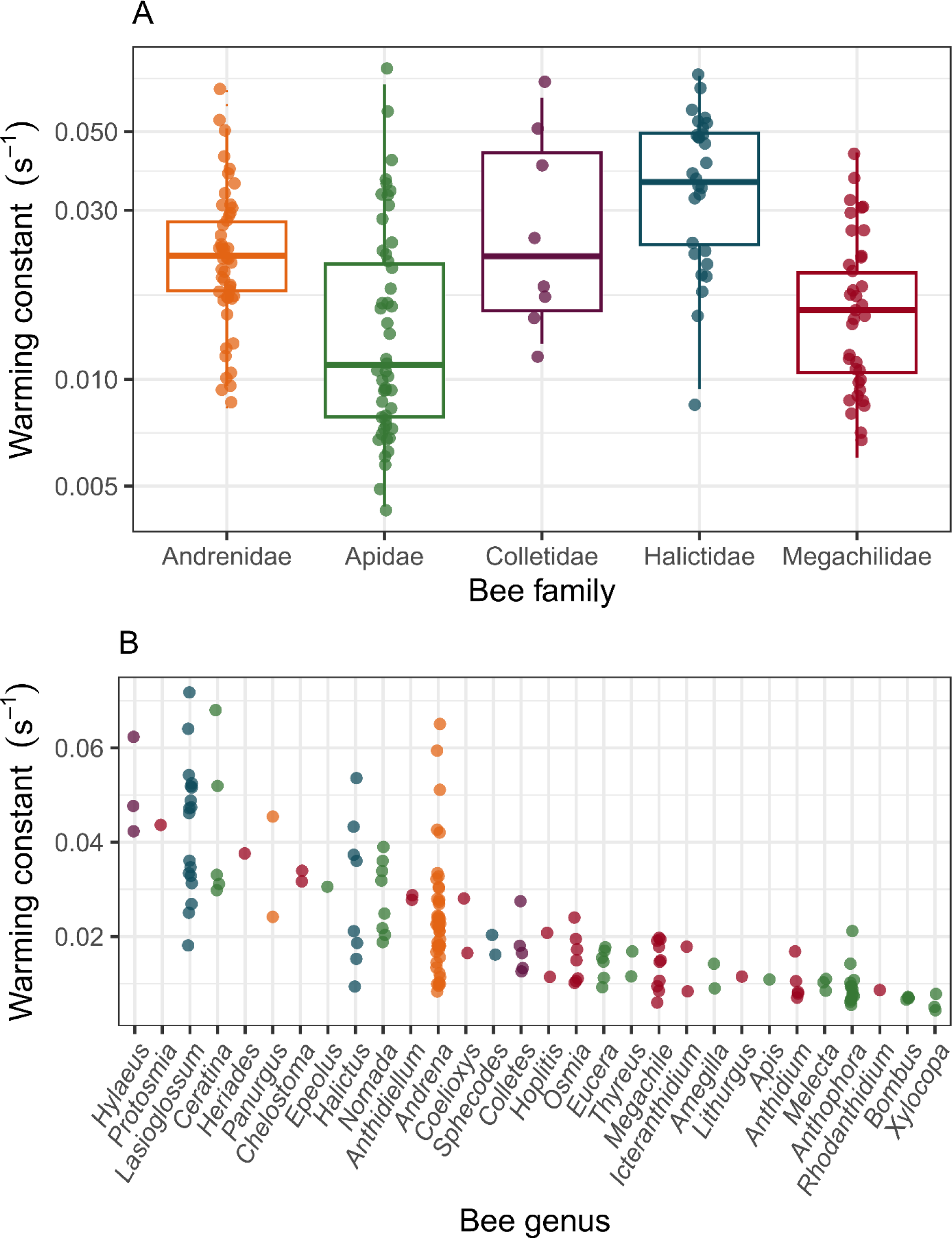
**A**. Mean, interquartile range and standard deviation of species means for warming constant (*K*) in the set of 169 bee species studied experimentally, shown separately by family. **B**. Mean warming constant (*K*) for individual species in the 30 genera represented in the species sample studied, depicted using the color codes for families in **A**. Genera are ranked from left to right in decreasing order of the average of species mean*s*. In both graphs, each symbol corresponds to an individual species and small random deviations were added to the data to improve readability. See Appendix S1: Tables S1 and S2 for list of species and sample sizes. Note logarithmic scale on vertical axes.

**Figure 2.**
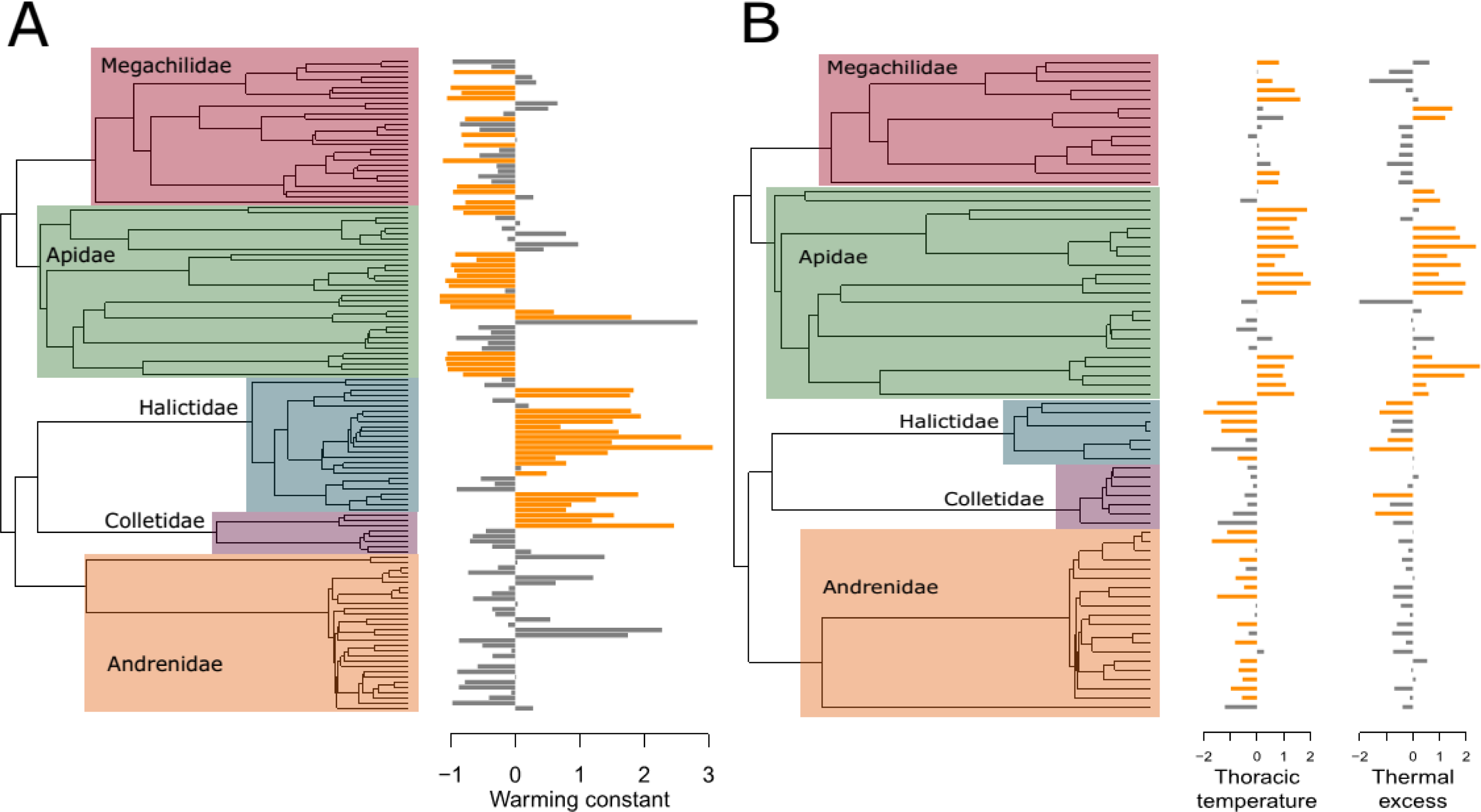
Visualization of species means for (**A**) warming constant (*K*), and (**B**) thoracic temperature (*T*_th_) and thermal excess (*T*_exc_) in relation to the phylogenetic trees depicting their evolutionary relationships for two subsets of the bee species studied. Tree size was determined in each case by availability of phylogenetic data in the most recent phylogenetic supertree of world bees (*N* = 125 and 71 species in **A** and **B**, respectively). Each horizontal bar corresponds to one bee species, and its length and direction denote its mean thermal parameter relative to the overall zero-centered sample mean, i.e., left- and right-facing bars correspond to values that are smaller and larger than the overall sample mean, respectively. Shown in orange are species with statistically significant local Moran’s *I*_i_ (*P* < 0.05), denoting instances of local phylogenetic associations. Clusters of orange-colored bars are indicative of “local hotspots” of phylogenetic autocorrelation for the thermal parameter involved. The monophyletic clades corresponding to the five bee families represented in this study are indicated in each graph by overlaid rectangles.

### Bee thermal biology: thoracic and ambient temperatures

Bees sampled in the field were foraging over an extremely broad range of thermal microenvironments (*T*_air_ range = 5.1–40.6°C, interquartile range = 18.2–27.0°C), and also exhibited a very broad range of thoracic temperatures (*T*_th_ range = 20.1–45.6°C, interquartile range = 29.5–38.7°C). When individual measurement of *T*_th_ and *T*_air_ (all bee species combined) were plotted on the corresponding bivariate space (Figure 3), data points exhibited two distinct patterns. Firstly, a conspicuously triangular distribution that virtually filled the space’s region satisfying the condition *T*_th_ > *T*_air_, thus revealing a consistently positive thermal excess of bees relative to the environment; extensive variability of such thermal excess across the range of environmental temperature; and a trend of thermal excess to decline with increasing ambient temperature. And secondly, there were two clusters of data points along the *T*_th_ axis that remained discernible over the whole range of variation in *T*_air_ (Figure 3). In contrast with the unimodal marginal distribution of *T*_air_ values (*D* = 0.00794, *P* = 0.63, Hartigans’ dip test), the marginal distribution of *T*_th_ values was significantly bimodal (*D* = 0.02015, *P* = 9.0E-06). As shown in the next paragraph, such clustering mostly reflected the distinctness of bee families and genera with regard to the particular thermal subspace with which they were predominantly associated. Species means for *T*_th_, *T*_air_ and *T*_exc_ were all significantly related to family and genus after statistically accounting for the (significant) effect of body mass (Table 1), thus denoting a taxonomic component to variation in thermal parameters. Irrespective of family or genus, however, larger bees tended to have higher *T*_th_ and *T*_exc_, and occurred at lower ambient temperatures, than smaller ones (Table 1).

**Figure 3.**
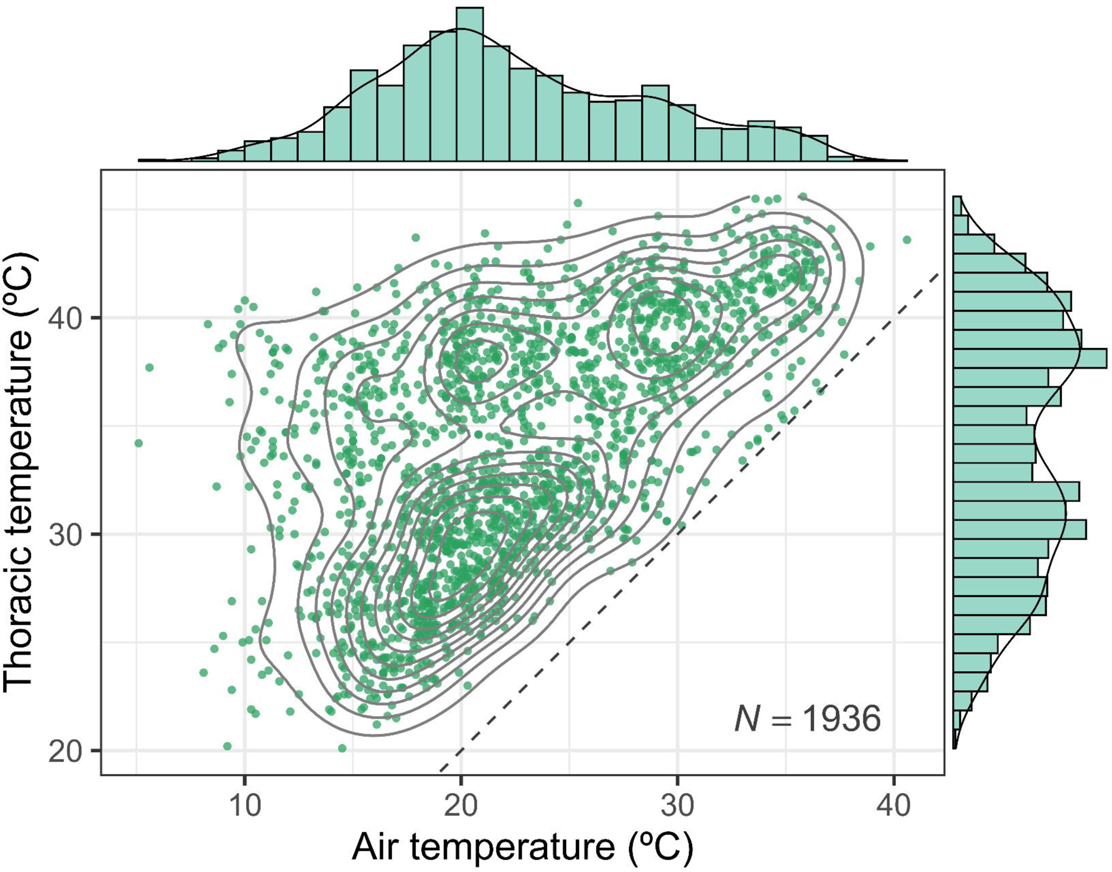
Bivariate distribution of paired measurements taken in the field of thoracic temperature of foraging bees (*T*_th_) and air temperature at capture spots (*T*_air_), all species combined. Contour lines depict the fitted two-dimensional kernel estimation of data point density; top and side marginal distributions represent histograms and univariate kernel-estimated distributions of *T*_air_ and *T*_th_, respectively; *N* = total sample size; and the dashed line depicts the *y* = *x* isothermal line.

Each bee family tended to occupy distinct regions in the plane defined by *T*_th_ and *T*_air_ (Figure 4). Family means for *T*_th_ fell into either “cool-” or “warm-bodied” groups, namely Halictidae (26.1 ± 0.44°C), Andrenidae (28.1°C ± 0.12°C) and Colletidae (29.9 ± 0.22°C) on one side, and Megachilidae (36.0 ± 0.20°C) and Apidae (37.8 ± 0.13°C) on the other, respectively (Figure 4). Mean thoracic temperatures of the coolest (Halictidae) and warmest (Apidae) families encompassed a range of ∼11°C. The ranking of bee families with regard to mean *T*_exc_ did not exactly match their *T*_th_ ranking, but they likewise tended to segregate into the same cool-(Halictidae, Andrenidae, Colletidae) and warm-bodied (Apidae, Megachilidae) groups with regard to *T*_exc_ (Figure 4). In some cases genera in the same family differed in the pattern of covariation between *T*_th_ and *T*_air_, but these differences were generally small in comparison with differences among families (Figure 5). Within the Apidae, the family with most genera in my sample, the genera *Amegilla*, *Anthophora*, *Apis*, *Bombus* and *Xylocopa* had similarly high *T*_th_ and *T*_exc_ values, while these were low in *Ceratina* and intermediate in *Eucera*. All genera of Megachilidae had similarly high *T*_th_ and *T*_exc_. On the opposite extreme, the two genera of Halictidae (*Halictus*, *Lasioglossum*) were similar in having low *T*_th_ and *T*_exc_ (Figure 5).

**Figure 4.**
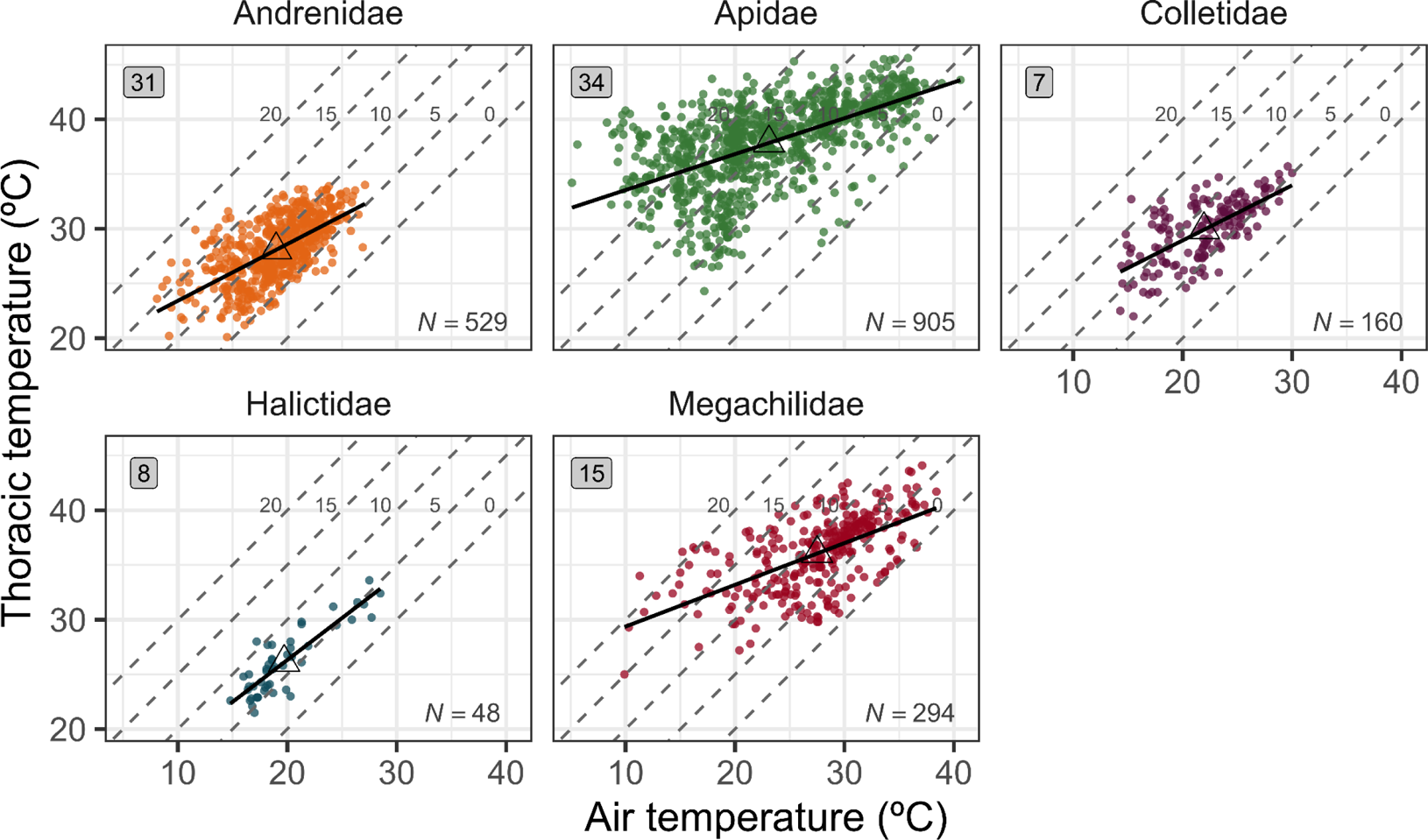
Bivariate distributions of field measurements of thoracic temperature of foraging bees (*T*_th_) and air temperature at capture spots (*T*_air_) separately for the different families. Individual graphs show raw measurements (dots), least-squares fitted regression (black line), bivariate mean (open triangles), and number of data points (*N*, bottom right) and species (rounded square, top left) included in the graph. Diagonal dashed lines demarcate areas with different levels of *T*_exc_ on the plane defined by *T*_th_ and *T*_air_, running from 0°C to +20°C from right to left (*y* = *x*, *y* = *x* + 5, *y* = *x* + 10, *y* = *x* + 15 and *y* = *x* + 20, respectively).

**Figure 5.**
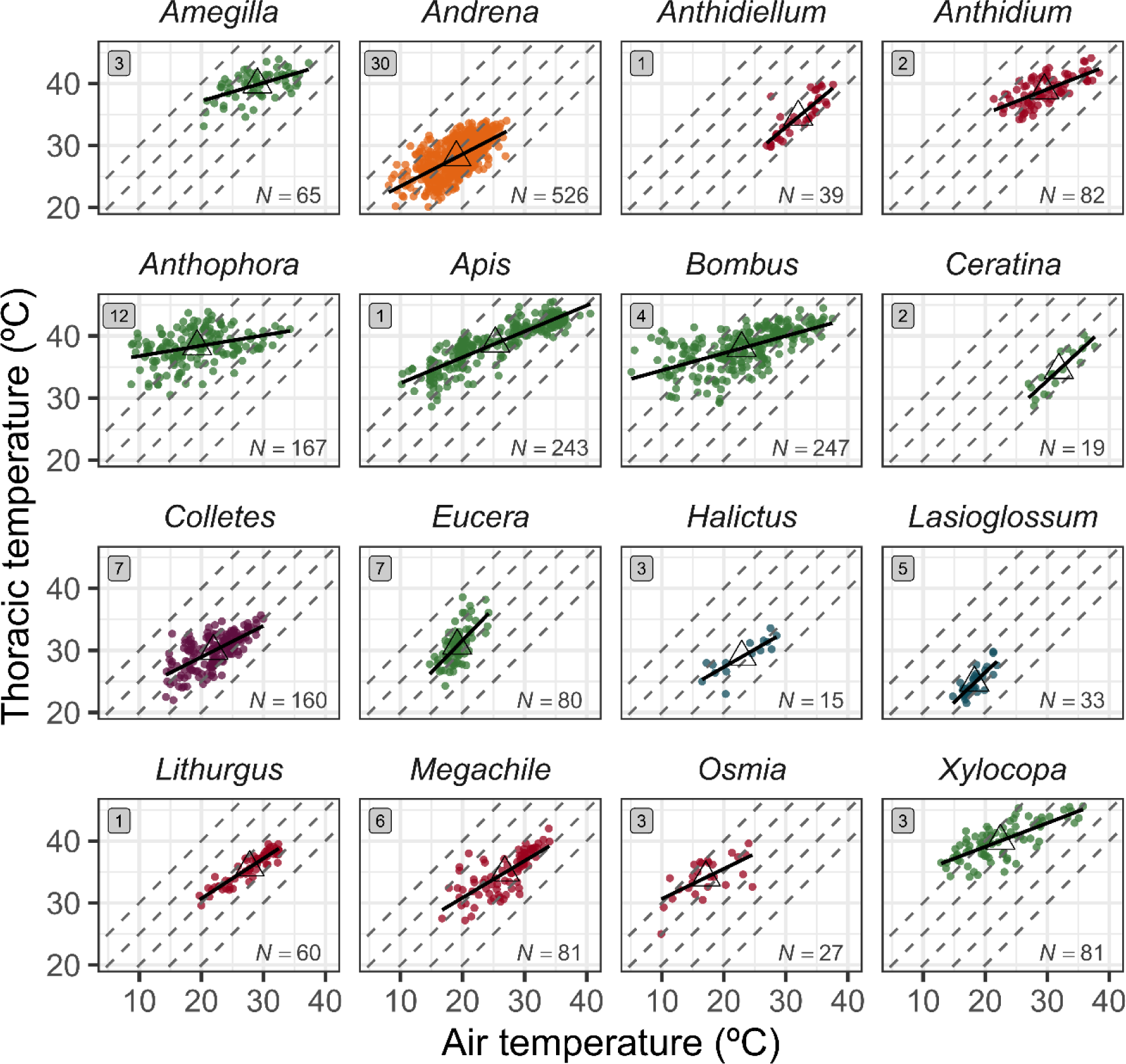
Bivariate distributions of field measurements of thoracic temperature of foraging bees (*T*_th_) and air temperature at the capture spots (*T*_air_) separately for the 16 genera having at least *N* = 15 data points. Genera are arranged alphabetically and color-coded by family as in Figure 4. Individual graphs show raw measurements (dots), least-squares fitted regression (black line), bivariate mean (open triangles), and number of data points (*N*, bottom right) and species (rounded square, top left) included in the graph. Diagonal dashed lines demarcate areas with different levels of *T*_exc_ on the plane defined by *T*_th_ and *T*_air_, running from 0°C to +20°C from right to left as labelled in Figure 4 (*y* = *x*, *y* = *x* + 5, *y* = *x* + 10, *y* = *x* + 15 and *y* = *x* + 20, respectively).

Heterogeneity among bee families, and among genera within families, in mean *T*_th_ and mean *T*_exc_ showed strong phylogenetic signals, as denoted by large and highly significant Pagel’s λ tests (Table 2), and distinctly clustered distribution of local Moran’s *I* values across the phylogenetic tree of species represented in the sample (Figure 2B).

**TABLE 2.** Tests for the presence of phylogenetic signal in species means for warming constant (*K*), thoracic temperature (*T*_th_) and thermal excess (*T*_exc_ = *T*_th_ – *T*_air_).

| Thermal biology<br>parameter | Species in<br>phylogenetic tree | Pagel's $\lambda$ | |
| --- | --- | --- | --- |
| | | Statistic | $P$ * |
| $K$ | 125 | 0.9196 | <1E-06 |
| $T_{\text{th}}$ | 71 | 0.8343 | <1E-06 |
| $T_{\text{exc}}$ | 71 | 0.9539 | <1E-06 |

Notes: \*, $P$ -value obtained by permutation with $10^6$ repetitions.

### The bee pollinator assemblage

Bees were the numerically dominant pollinators of entomophilous plants in the Sierra de Cazorla area. They were recorded in the pollinator censuses of 95.9% of the plant species studied (*N* = 292), and accounted for 44.4% of all pollinator individuals (*N* = 41,948) and 59.0% of all flower visits (*N* = 127,493) recorded, all pollinator and plant species combined. The regional bee pollinator assemblage was extraordinarily species-rich. A total of 250 species in 37 genera were identified in censuses, roughly comprising 60% of all bee species reported to date for the region (Ortiz-Sánchez et al. 2023). The family Apidae contributed 36.9% of all bee individuals (*N* = 18,622), followed in decreasing order of importance by Halictidae (24.9%), Megachilidae (16.1%), Andrenidae (14.2%) and Colletidae (7.8%). Most bee individuals belonged to the genera *Lasioglossum*, *Bombus, Andrena*, *Halictus*, *Ceratina*, *Hylaeus* and *Heriades*, mentioned in decreasing order of frequency (Figure 6). These seven genera accounted altogether for 67.5% of bees recorded in censuses.

**Figure 6.**
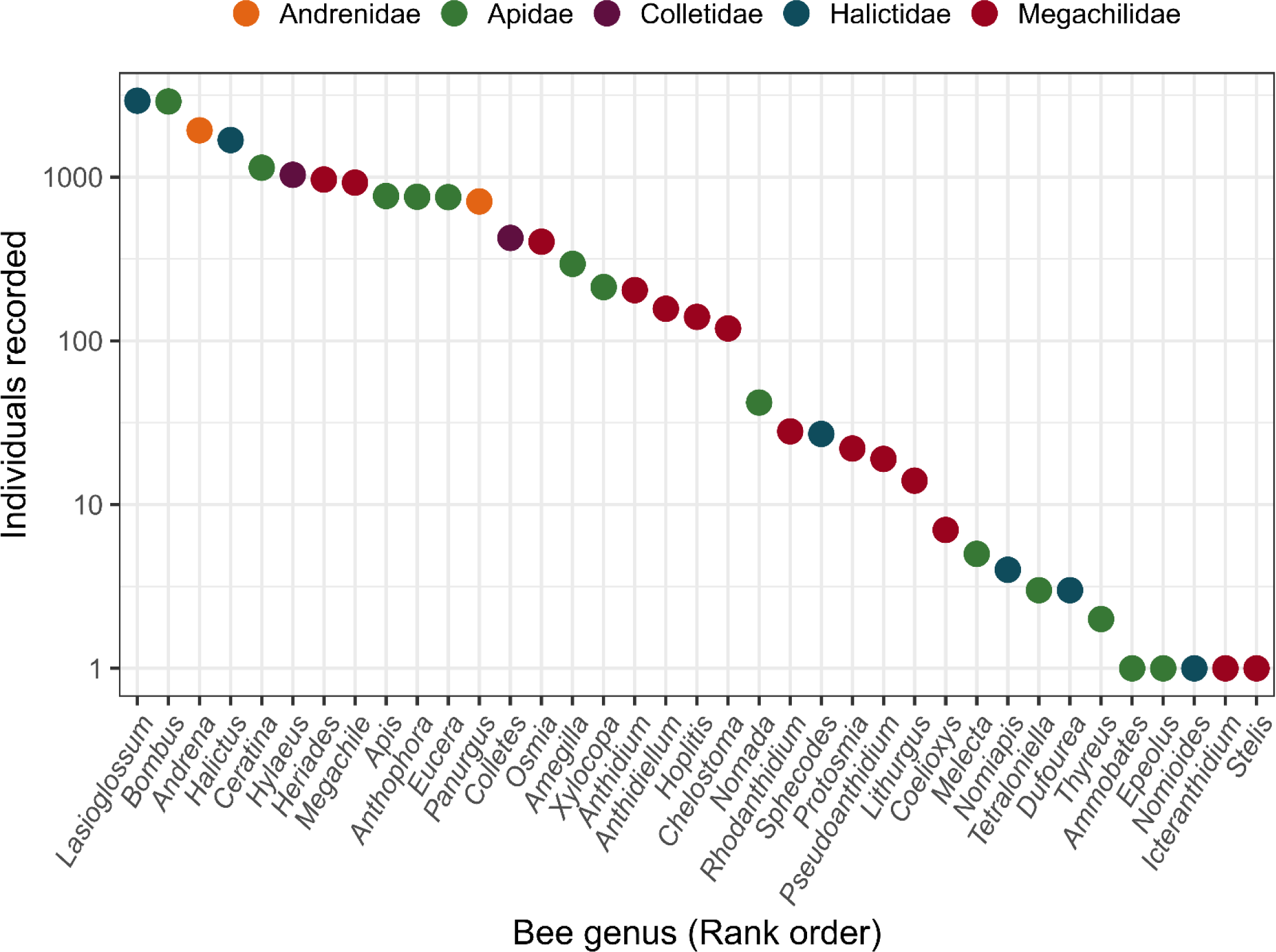
Genus abundance curve (ranked number of individuals per genus) for bees recorded in pollinator censuses conducted on a total of 292 entomophilous plant species from the Sierra de Cazorla area, southeastern Iberian Peninsula, all plant species combined (*N* = 18,622 bee individuals). Note logarithmic scale on vertical axis.

### Temporal patterns: seasonal and daily variation in thermal profile

Community-wide inferences on bee thermal parameters (*K*, *T*_th_, *T*_exc_), obtained by assigning species means to the bee individuals recorded in censuses, revealed significant seasonal trends in the warming constant, thoracic temperature and thermal excess of the bee assemblages visiting the flowers of the plant community studied (Figure 7). Bees with low warming constants (< 0.04 s^-1^) prevailed during March-May, while individuals from species with higher values (0.04-0.07 s^-1^) joined the assemblage during June-August, which contributed to produce a marked seasonality (Figure 7A). Thoracic temperature and thermal excess of individual bees both tended to rise from March through May, but while *T*_th_ remained stable afterwards, *T*_exc_ declined steadily through September, largely reflecting the disappearance from the bee community during the summer of species with typically large *T*_exc_ values (Figure 7B, 7C). The preceding descriptions of seasonal patterns, based only on the inspection of fitted nonparametric regressions plotted in Figure 7, were upheld by analytical results obtained by running three generalized additive models each with one thermal parameter as response variable, a predictor smooth term for Julian date (days from 1 January), and log body mass, bee family, and bee genus (nested within family) as parametric fixed-effect covariates to statistically account for their effects (Appendix S1: Table S4).

**Figure 7.**
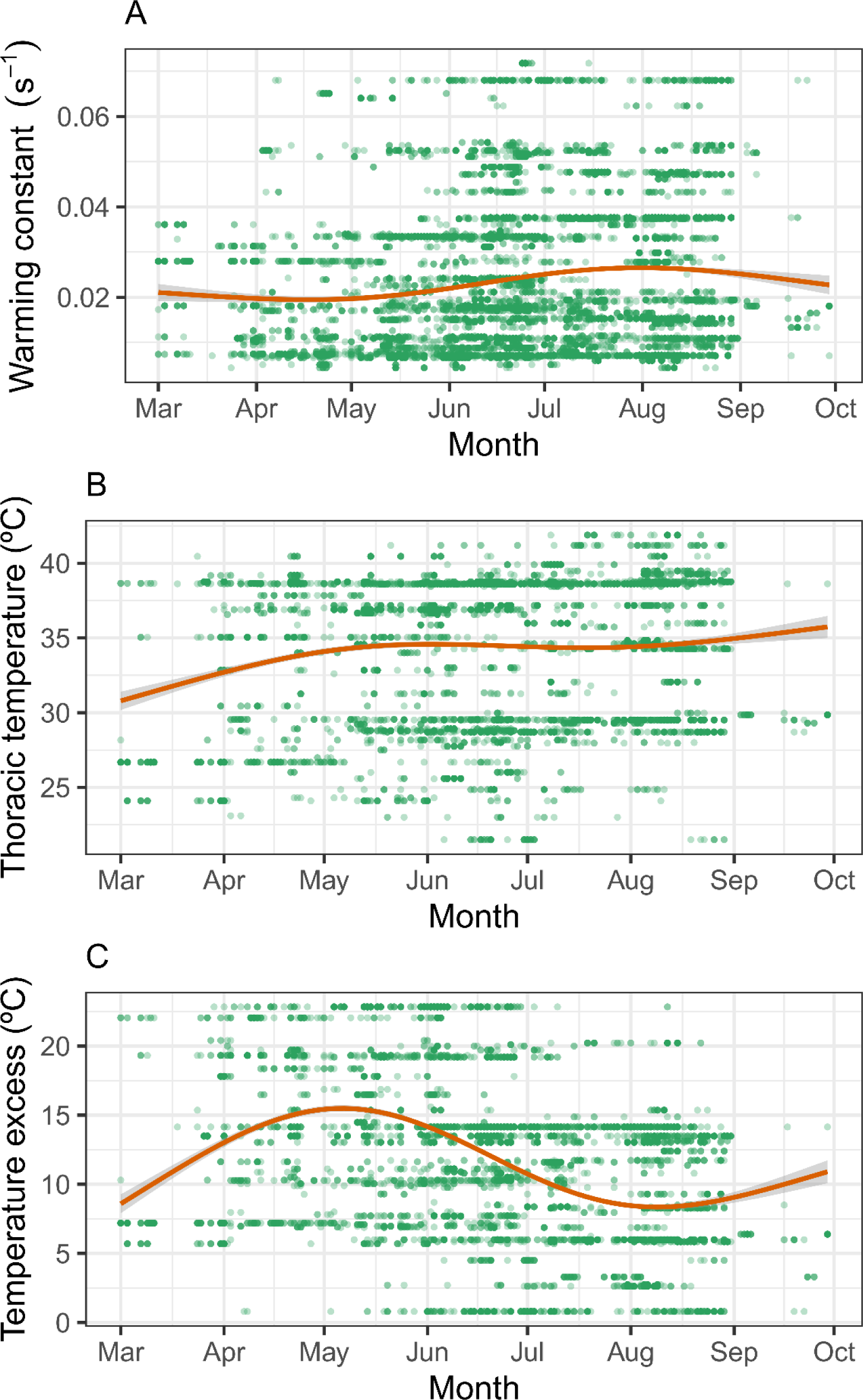
Variation over March-September in (**A**) warming constant (*K*), (**B**) thoracic temperature (*T*_th_) and (**C**) thermal excess (*T*_exc_) inferred for individual bees identified to species in pollinator censuses. Each dot represents a bee belonging to a species with *K*, *T*_th_, *T*_exc_ means available to be associated with individual bee observations (*N* = 14169, 8336 and 8336, for *K*, *T*_th_ and *T*_exc_, respectively). Red lines represent nonparametric regressions fitted to the data using a smoothing spline and their associated 95% confidence intervals (grey bands).

Patterns of daily variation in thermal parameters (*K*, *T*_th_, *T*_exc_) of the bee assemblages recorded in pollinator censuses were examined separately by month, since seasonal rhythms in time of sunrise, daytime length and ambient temperature could induce seasonal changes in daily patterns. Results are shown in Figure 8. Irrespective of time of year, there was a consistent trend for the warming constant of bees to be lowest among those flying early in the morning and then increasing steadily up to reaching a maximum or plateau around or shortly after noon (Figure 8A). This daily pattern was mainly a consequence of high-*K* bees being largely restricted to the central hours of daytime. Daily variation in thoracic temperature and thermal excess roughly mirrored the pattern for *K*. Both *T*_th_ and *T*_exc_ tended to decline from early morning to noon, except in May and June when these parameters remained fairly constant from sunrise to noon (Figure 8B, 8C).

**Figure 8.**
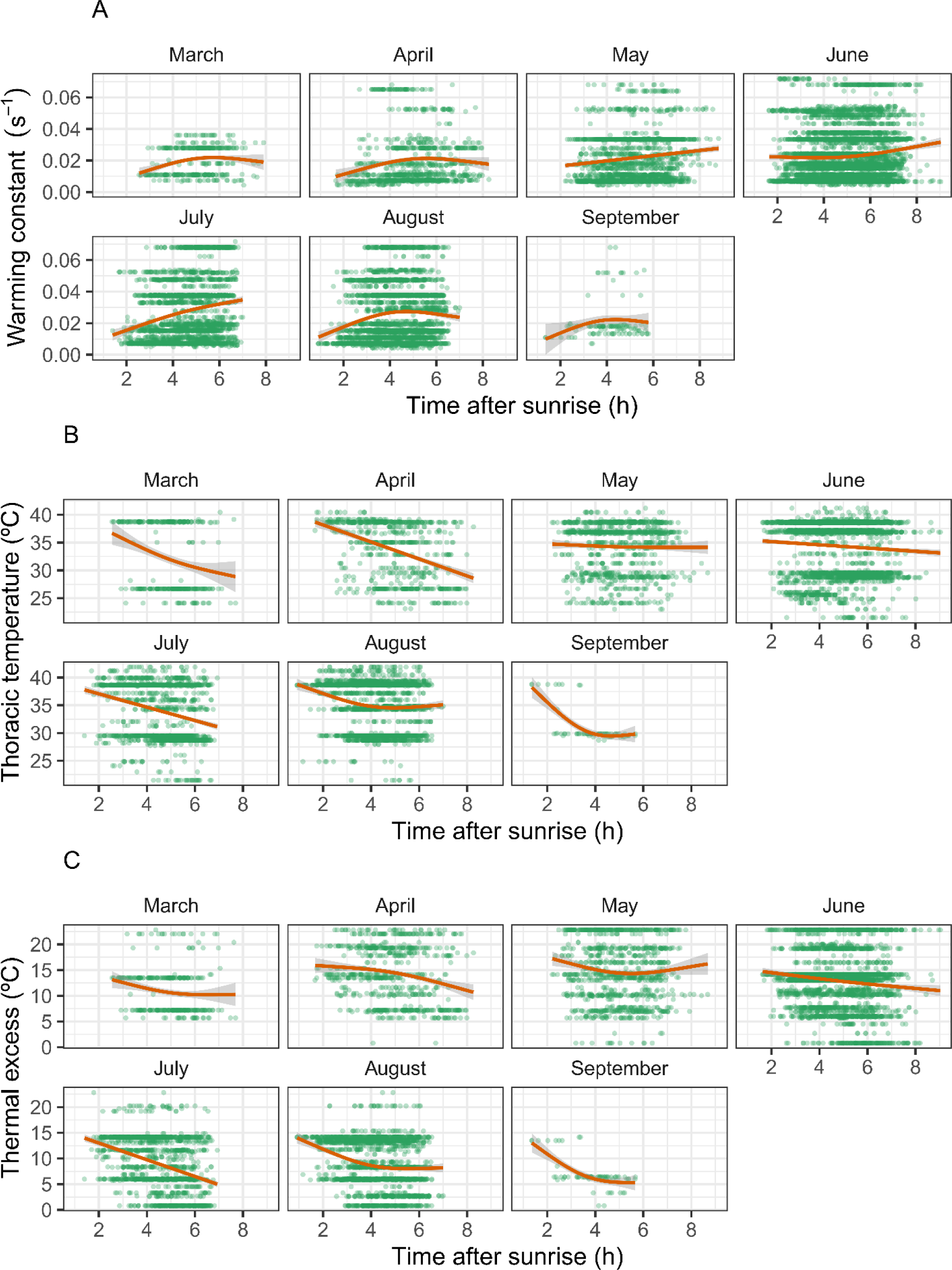
Daily variation, and its monthly fluctuations over the period March-September, in (**A**) warming constant (*K*), (**B**) thoracic temperature (*T*_th_) and (**C**) thermal excess (*T*_exc_), inferred for individual bees which were identified to species in pollinator censuses. Each dot represents a bee belonging to a species with *K*, *T*_th_, *T*_exc_ means available to be associated with individual bee observations (sample sizes as in Figure 7). Red lines depict nonparametric regressions fitted to the data using a smoothing spline and their associated 95% confidence intervals (grey bands).

### Spatial patterns: among-habitat variation in thermal profile

The bee pollinator assemblages of plants from different habitat types were significantly heterogeneous with regard to *K* (Chi-squared = 808.2, df = 8, *P* < 2.2E-16, Kruskal-Wallis rank sum test), *T*_th_ (Chi-squared = 417.2, df = 8, *P* < 2.2E-16) and *T*_exc_ distributions (Chi-squared = 730.1, df = 8, *P* < 2.2E-16) of individuals recorded, all plant species in each habitat type combined (see sample sizes separately by habitat type in Appendix S1: Table S5). For example, mean warming constant tended to be lowest in the bee pollinator assemblages of plants from tall sclerophyllous scrub, rock cliffs and forest interior; average thoracic temperatures were highest for bees visiting the flowers of plants from tall sclerophyllous scrub, forest interior and dolomitic outcrops; and thermal excess was lowest at streams/springs, grasslands/meadows and disturbances, and highest at cliffs and forest interior (Figure 9). Heterogeneity among habitat types in bee thermal parameters remained statistically significant when it was tested by means of linear models in which bee body mass, family and genus were included as covariates (Appendix S1: Table S6), thus denoting that among-habitat differences in bee thermal profiles were not entirely due to major differences in either taxonomic composition or body size of the respective bee assemblages.

**Figure 9.**
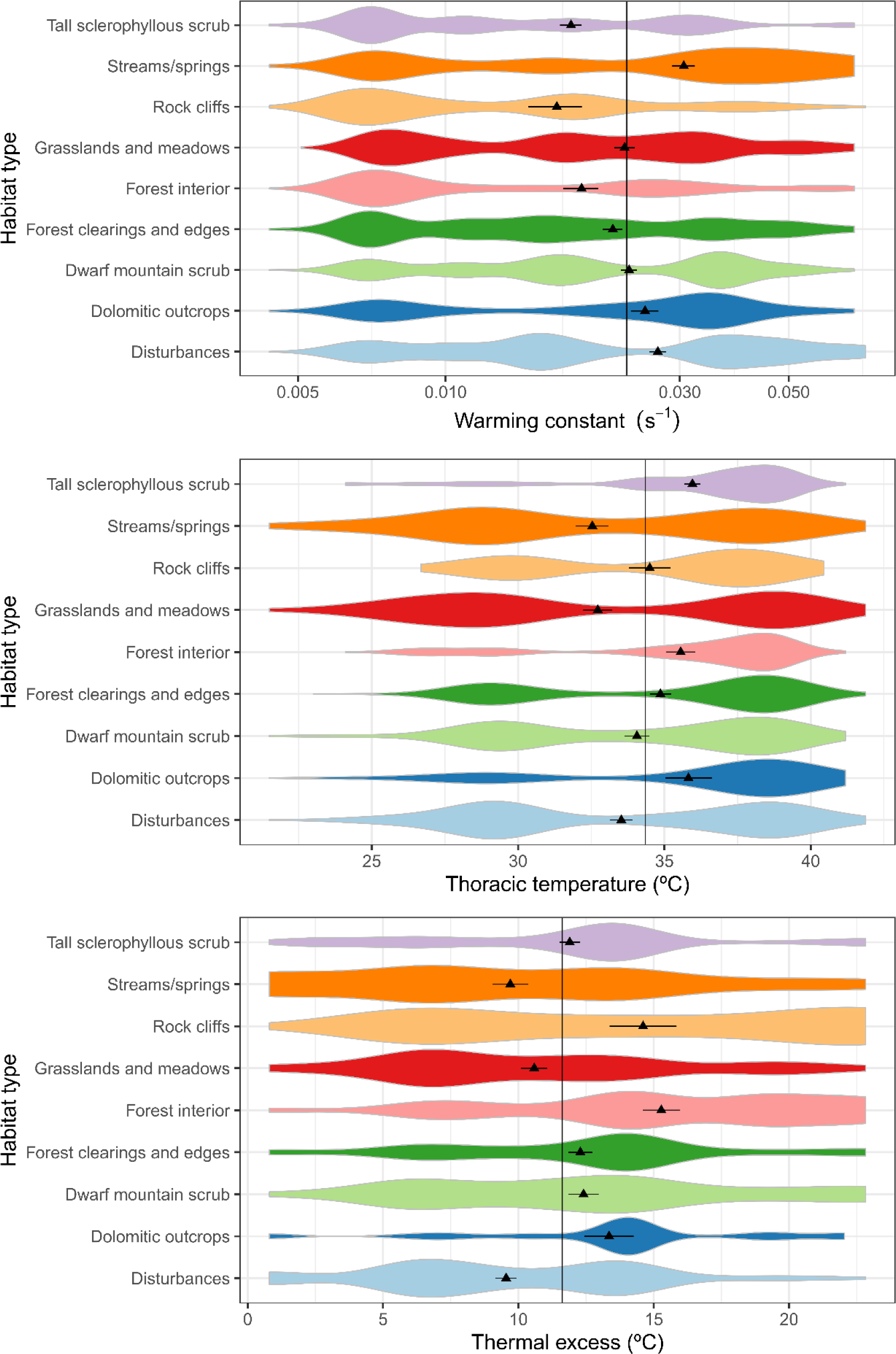
Inferred distributions of warming constant (*K*), thoracic temperature (*T*_th_) and thermal excess (*T*_exc_) in the bee pollinator assemblages of plant species from different habitat types (all plant species combined within each habitat type). Inferred distributions were constructed by assigning to each individual bee recorded in censuses and identified to species the corresponding species mean for *K*, *T*_th_ and *T*_exc_. Triangles denote averages (± 3SE) and the thin black vertical line is the mean value for all habitat types combined, shown for reference. Note horizontal logarithmic scale in the uppermost graph. See Appendix S1: Table S5 for sample sizes (number of data points and species included in each distribution), and Appendix S1: Table S6 for results of statistical analyses.

Heterogeneity in central trends, however, did not completely capture the nature of differences among habitat types in the thermal biology profile of bee assemblages, as illustrated by kernel estimates of distributions represented as violin plots in Figure 9. There was an overall trend towards bimodal (*T*_th_, *T*_exc_), or sometimes trimodal (*K*) distributions, denoting that bee individuals tended to group into distinct clusters falling on particular segments of the thermal parameters’ axes. The relative importance of particular clusters differed among habitats. For example, “warm-bodied” bees with low *K* and/or high *T*_th_ and *T*_exc_ predominated in tall sclerophyllous scrub, rock cliffs, forest interior and dolomitic outcrops, while more even mixtures of different thermal types were found in the rest of habitat types (Figure 9).

### Bee thermal biology and flower visitation rate

Each of the three parameters used to characterize thermal biology of bees (*K*, *T*_th_, *T*_exc_) was a significant predictor of flower visitation rate (= number of flowers visited by each bee per 3-min watching period) (Table 3). These effects occurred independently of variations in the number of flowers available in the census patch and the bee species’ body mass, both of which were included as covariates in the linear mixed models and had positive effects on visitation rates (more flowers were visited by time unit in patches with more flowers and/or by larger bees; Table 3). The inclusion of plant species as a random effect in the models ensured that these conclusions were robust to interspecific variation in overall bee visitation rates or other plant-specific factor not accounted for here.

**TABLE 3.** Summary of linear mixed-effect models testing for the effects of corolla type (open *vs.* restrictive), individual bees’ inferred thermal parameters (*K*, *T*_th_ and *T*_exc_), and their interaction, on flower visitation rate (number of flowers visited per 3-min, log-transformed). Total number of flowers available in the patch and the individual bees’ inferred body mass were included as covariates. Plant species was included as a random effect in all models (results not shown).

| Thermal parameter tested for effect on flower visitation (sample sizes) | Predictor (fixed effects) | Parameter estimate ± SE | Chi-squared | *P*-value |
| --- | --- | --- | --- | --- |
| Warming constant ($K$)<br>($N = 14921$ bees, 272 plant species) | $K$ | $0.32 \pm 0.53$ | 10.1 | 0.0015 |
|  | Corolla type (CT) |  | 1.3 | 0.26 |
| | $K \times CT$ | | 33.7 | 6.3E-09 |
| | Flowers in patch* | $0.27 \pm 0.01$ | 633.1 | <2.2E-16 |
| | Bee body mass* | $0.15 \pm 0.01$ | 120.8 | <2.2E-16 |
| Thoracic temperature ($T_{th}$)<br>($N = 8951$ bees, 248 plant species) | $T_{th}$ | $0.00019 \pm 0.002$ | 14.1 | 0.00017 |
|  | Corolla type (CT) |  | 0.3 | 0.58 |
| | $T_{th} \times CT$ | | 10.1 | 0.0015 |
| | Flowers in patch* | $0.30 \pm 0.01$ | 469.3 | <2.2E-16 |
| | Bee body mass* | $0.13 \pm 0.01$ | 115.0 | <2.2E-16 |
| Thermal excess ($T_{exc}$)<br>($N = 8951$ bees, 248 plant species) | $T_{exc}$ | $0.0032 \pm 0.002$ | 44.0 | 3.3E-11 |
|  | Corolla type (CT) |  | 0.26 | 0.61 |
| | $T_{exc} \times CT$ | | 7.42 | 0.0064 |
| | Flowers in patch* | $0.30 \pm 0.01$ | 474.0 | <2.2E-16 |
| | Bee body mass* | $0.11 \pm 0.01$ | 76.0 | <2.2E-16 |

Notes: Predictors marked with asterisks were log-transformed for the analyses.

The main effects of bee thermal parameters on flower visitation rate were interpretable only in relation to corolla type (open vs. restrictive), given the statistically significant interactions between this variable and each thermal parameter (Table 3). Flowers of the two types differed markedly with regard to the slope of the relationship between visitation rate and bee thermal parameter, as illustrated by interaction plots of marginal effects (i.e., holding constant the effects of number of flowers in the watched patch and bee body mass). In plant species with restrictive corollas there was a steep decline of flower visitation rate with increasing warming constant, and steep increases with increasing thoracic temperature and thermal excess (Figure 10). In contrast, the relationship between flower visitation and thermal parameters was weak (*K*, *T*_exc_) or null (*T*_th_) in plants with open corollas (Figure 10). As a consequence of these interactions effects, flowers with open corollas were at a bee visitation advantage over flowers with restricted corollas when they were visited by bees of high *K* (>∼0.02 s-^1^) and low *T*_th_ (< ∼30°C) and *T*_exc_ (<∼8°C), and vice versa (Figure 10).

**Figure 10.**
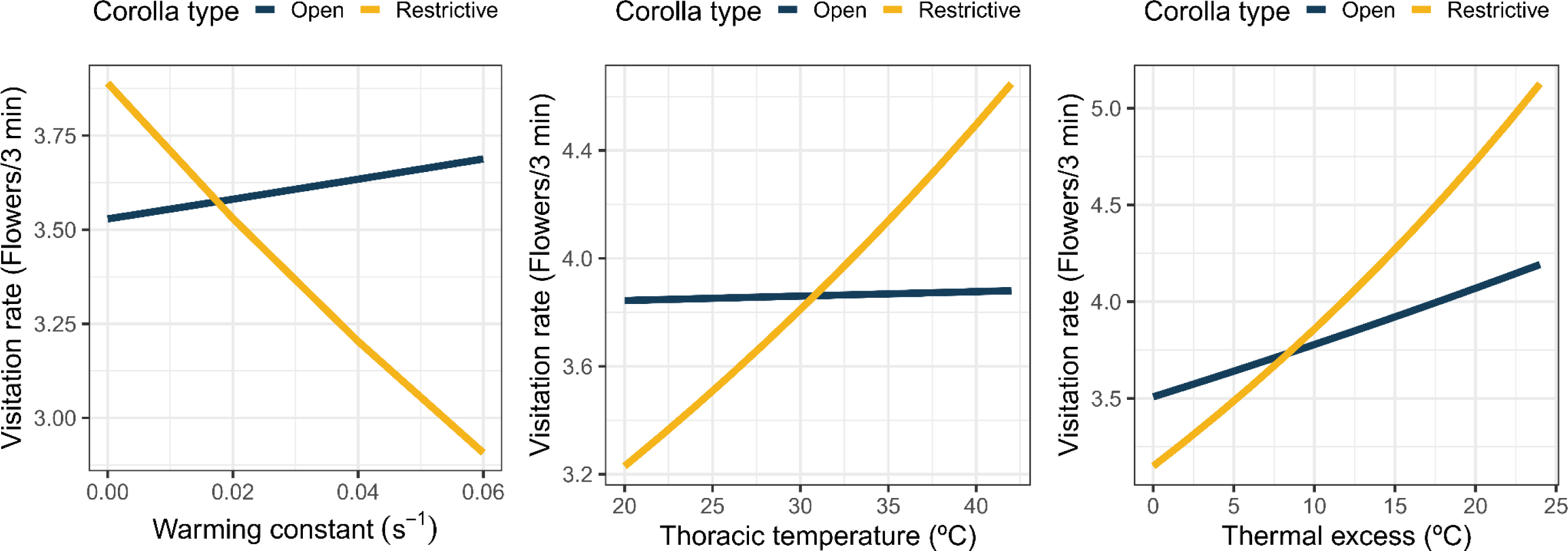
Interaction plots depicting the marginal effects (i.e., holding constant the effects of bee body mass and number of flowers in the watched patch; Table 3) of thermal parameters (*K*, *T*_th_, *T*_exc_, from left to right) on the flower visitation rate of individual bees in plants with open and restrictive corollas. See Table 3 for a summary of the three corresponding linear mixed-effect models (plant species was included as a random effect in every model), including the statistical significance levels for the three corolla type x thermal parameter interactions illustrated here.

## DISCUSSION

By examining a sizeable fraction of the diverse bee pollinator assemblage associated with the community of entomophilous plants from the Sierra de Cazorla area, this study found that these key pollinators harbor considerable diversity with regard to thermal biology features. This diversity largely stems from the high taxonomic diversity at the family, genus and species levels represented in my sample, since different taxonomic groups have distinct thermal features. Taxonomy-related diversity of thermal features is, in turn, associated with a strong phylogenetic signal that is mostly expressed at the level of bee families, which suggests an early origin of thermal diversification in the evolutionary history of bees. Furthermore, bee pollinator assemblages associated with different habitats, time of day or time of year tended to differ in their thermal parameters. This lends support to the hypothesis formulated in the Introduction that diversity in thermal biology of bees can eventually enhance opportunities for bee coexistence, spatiotemporal partitioning of floral resources, and pollination success for plants, as discussed below. Sampling thoroughness of the regional communities of both plants and bees confers significant robustness to the conclusions of this study.

### Thermal biology diversity

Intrinsic heat transfer rate (*K*), temperature of thoracic flight muscles while foraging at flowers (*T*_th_), air temperature at foraging site (*T*_air_), and thermal excess (*T*_exc_) of individual bees have been used here to characterize their thermal biology. This set of parameters represents a rather minimalist subset of the possible thermal features that could have been considered. Aspects not considered include, for instance, endothermic and thermoregulatory ability, or the relationship between thoracic and operative temperatures, previously addressed in comparative studies of bee thermal biology (Stone 1993a,b, Stone and Willmer 1989, Herrera 1995, Bishop and Armbruster 1999, Herrera et al. 2023). Inclusion of these aspects might have produced a higher-resolution picture of thermal biology diversity and their ecological correlates, but this improvement at resolution would come at the cost of reducing the number of taxa examined, since there was a compromise between number of taxa and the thoroughness of thermal information obtainable per taxon. Since the main focus of this study was evaluating thermal diversity and its taxonomic, phylogenetic and ecological correlates, collecting data for many species was essential for obtaining a robust, well-populated dataset after merging thermal information with the data from pollinator censuses. Depth of thermal information for individual species was therefore sacrificed in favor of sampling a larger number of species. The thermal parameters used, however, proved sufficient to document the thermal diversity of the regional bee pollinator assemblage and its correlates.

Laboratory estimates of warming constant (*K*) revealed extensive variation of this parameter in the bee species sample considered here. The *K* value of an inert object estimates the rate at which it gains heat when placed in an environment warmer than itself, or losses heat in a cooler environment. *K* is inversely, nonlinearly related to body mass and, in the case of insects, depends also on inherent properties influencing heat transfer such as coat color, body geometry, reflectance, pubescence or evaporative water loss (May 1976, Willmer and Unwin 1981, May and Casey 1983, Bressin and Willmer 2000, Pereboom and Biesmeijer 2003, Herrera et al. 2023). The broad variation in *K* among bee taxa found here denoted intrinsic heterogeneity in the rate at which heat leaves or enters individual bees’ bodies when other influential factors (e.g., behavioral or physiological thermoregulation) were controlled experimentally by taking the measurements on dead bees. In bees, variation in heat gain/retention ability is known to correlate with the relative importance of endothermy vs. ectothermy, with predominantly endotherms having lower *K* values than predominantly ectotherm ones, and larger thermal excess in relation to the ambient (May 1976, Herrera et al. 2023). Results of this study likewise point to an inverse relationship across bee families, and across genera within families, between *K* values and thermal excess in the field (compare Figures 1, 4 and 5). *K* estimates could thus be profitably used as meaningful proxies for assessing thermal biology diversity in large samples of bee species, as they can be obtained for bees of all sizes and require less effort than the very time-consuming field measurements of *T*_th_ and *T*_air_.

Diversity of thermal biology had a strong taxonomic basis. The long-known relationships between body mass and thermal biology parameters in insects (e.g., Digby 1955, May 1976, Unwin and Corbet 1984, Stone and Willmer 1989, Stone 1993a,b, Bishop and Armbruster 1999, Rodríguez et al. 2018, Herrera et al. 2023) was also consistently found in this study. Body mass was inversely related to *K* and directly to *T*_th_ and *T*_exc_ (Table 1). To statistically account for these effects, body mass was included as a covariate in all linear, linear mixed-effect, and generalized additive models fitted to the data (Tables 1 and 2, Appendix S1: Tables S4 and S6) to ensure that the effects of the predictors of interest were not concealed or distorted by the broad variation in body mass represented in the sample. After accounting for body mass variation, heterogeneity among bee families, and among genera nested within families, in thermal biology parameters emerged as a powerful force in structuring community-wide patterns of bee thermal biology. This conclusion strenghtens and extends those of previous studies based on smaller or taxonomically narrower species sets (Stone and Willmer 1989, Bishop and Armbruster 1999, Herrera et al. 2023). Species of Colletidae, Halictidae and Andrenidae, three major bee families which have been historically underrepresented in studies of bee thermal biology despite their abundance and diversity in regional bee communities worldwide (J. Herrera 1988, Kato 2000, Torné-Noguera et al. 2014, Turley et al. 2022), are well represented in my sample and contributed substantially to broaden the thermal biology space of the whole bee assemblage (Figure 4) (see Stone and Willmer 1989, Shelly et al. 1993, Herrera 1995, Potts 1995, Bishop and Armbruster 1999, Herrera et al. 2023, for earlier data on these understudied bee families).

Recent molecular phylogenies of bees show that all families and most genera considered in this study are monophyletic lineages (Bossert et al. 2022, Orr et al. 2022, Pisanti et al. 2022, Almeida et al. 2023, Henríquez-Piskulich et al. 2024), hence it is hardly surprising the finding of a strong phylogenetic signal underlying the taxonomic correlates of thermal biology diversity. Dated molecular phylogenies have documented a deep split occurring very early in the evolutionary history of bees (∼115 million years ago; Almeida et al. 2023, Henríquez-Piskulich et al. 2024) which led to two distinct lineages consisting of [Apidae + Megachilidae] on one side, and [Halictidae + Colletidae + Andrenidae] on the other. As illustrated in Figure 2, family-level differences in thermal biology parameters found in this study closely matched this major evolutionary split. In my species sample, taxa in the [Apidae + Megachilidae] clade were characterized by relatively low *K* and high *T*_th_ and *T*_exc_ (dubbed “warm-bodied” group above), while taxa in the clade formed by [Halictidae + Colletidae + Andrenidae] had a complementary combination of traits (the “cool-bodied” group). In addition, previous work indicated that endothermy tends to be more important than ectothermy in the “warm-bodied” group, while the reverse holds for the “cool-bodied” one (Herrera et al. 2023, and references therein), thus pointing to another divergent thermal trait associated with a major evolutionary split in the history of bees. The preceding observations therefore suggest that (1) bimodality found here in the distribution of bee temperatures in the face of unimodal ambient temperatures (Figure 3) is a thermal signature of an old, major divergence event in the evolutionary history of bees; (2) to a greater or lesser degree, evolutionary diversification of bees seems to have been associated with divergence in thermal biology; and (3) thermal diversification of bees may have contributed substantially to broadening the (ecological) breadth of thermal niches and floral resources exploited. Circumstantial evidence supporting these interpretations is provided by the correlated evolution of cell membrane composition with thoracic temperature and flight metabolic rate in bees (Rodríguez et al. 2015, 2018). Specifically, the relative abundances of palmitate, oleate and linoleate in membrane phospholipids are associated with family-level differences in thoracic temperature (Rodríguez et al. 2018). Since cell membrane composition is a crucial element in cell functionality (Hazel 1995) and thus expected to remain highly conserved evolutionarily, its early divergence among bee families could ultimately account for, and contribute to, extant thermal diversity.

### Ecological correlates

Daily variation in the thermal biology profile of bees tracked the natural course of temperature and solar irradiance from sunrise to noon, and each taxonomically-, phylogenetically- and thermally-defined major bee group (dubbed “warm-” and “cool-bodied” for simplicity here) tended to have a characteristic temporal window of activity. Regardless of time of year, bees with low *K* and high *T*_th_ and *T*_exc_ tended to prevail shortly after sunrise, the daytime period with the lowest incident solar irradiance and ambient temperature. This group of early-morning bees typically included species of large-sized Apidae in the genera *Bombus* and *Anthophora*, both of which are well known for their thermoregulatory abilities and strong endothermy (Heinrich 1993, Stone 1993a,b, 1994, Bishop and Armbruster 1999). Bees with progressively higher *K* and lower *T*_th_ and *T*_exc_ entered the bee assemblage as the morning proceeded, leading to the highest *K* and lowest *T*_th_ and *T*_exc_ averages being reached around noon. Although the taxonomic composition of this group of late-morning and midday bees tended to change from late winter-spring (mostly *Andrena* and *Lasioglossum*) to summer (mostly *Colletes*, *Halictus*, *Heriades*, *Hylaeus*), they consistently were small sized (as frequently reported by previous studies; e.g., Linsley 1978, Herrera 1990) and belonged to predominantly ectothermic genera and families (Andrenidae, Colletidae, Halictidae; Herrera et al. 2023, and references therein). In this way, low-*K* bee lineages (i.e., heat conservatives, putative endotherms) tended to be the main or sole exploiters of flowers during the cooler period of the day with low potential radiative heat gains, while the high-*K* ones (i.e., fast passive warming/cooling, putative ectotherms) tended to prevail during the warmest period of daytime, when intense solar radiation was available. It is therefore tempting to suggest that the early evolutionary split into low- and high-*K* lineages endowed bees as a group with an ability to successfully exploit flowers in both shady and sunny habitats.

Seasonal changes in thermal biology profiles were vaguely reminiscent of those taking place over daytime, since the increase in *K* from cooler spring to warmer summer bore some resemblance to daily variation in that the importance of cool-bodied bees was greater under warmer conditions. This was mainly brought about by increased representation in summer of the smallest sized species from each family (*Anthidiellum*, *Ceratina*, *Halictus* subgenera *Seladonia* and *Dialictus*, *Heriades*, *Hylaeus*), a seasonal pattern in body size already described for other Mediterranean bee communities (Shmida and Dukas 1990, Osorio-Canadas et al. 2016). The seasonal variation in *T*_th_ and *T*_exc_ of bee assemblages followed a more complex pattern, involving increases from early to late spring followed by either decline (*T*_exc_) or stabilization (*T*_th_) until the end of summer. These seasonal changes mainly reflected the virtual disappearance of cool-bodied, predominantly ectothermic *Andrena* by the end of spring (Herrera et al. 2023) and the increased representation throughout the summer of warm-bodied, putative endotherms in the Apidae (*Amegilla*, *Anthophora* subgen. *Heliophila*, and *Xylocopa*) and Megachilidae (mostly species of *Anthidium*, *Lithurgus* and *Megachile*) (C. M. Herrera 1988, 1990, 1997).

The composition and diversity of bee assemblages generally vary among habitats in the Mediterranean region (Nielsen et al. 2011, Torné-Noguera et al. 2014, Reverté et al. 2019, Gómez-Martínez et al. 2022, Pérez-Marcos et al. 2023). Results of this study show that, as a consequence of the taxonomic and phylogenetic correlates of thermal biology, such variation in composition eventually translates into habitat-specific species sets which have distinctive profiles of thermal biology. The nine habitat types recognized here differed in both central trend and spread of distributions of the bee thermal parameters considered. Predominantly shady habitats (forest interior, dense Mediterranean sclerophyllous scrub and rock cliffs) were characterized by thermally less diverse bee assemblages and prevalence of taxa with low *K* and high *T*_th_ and *T*_exc_. In contrast, bee assemblages from predominantly sunlit habitats (e.g., forest clearings and edges, grasslands and meadows) were characterized by the broadest variety of thermal biology features. These among-habitat differences can be parsimoniously interpreted in terms of the differential reliance of major bee groups on solar irradiance to attain the thoracic temperatures needed for flight. Only low-*K*, putatively endothermic taxa largely independent of solar irradiance (mostly large, fast-flying Apidae such as *Anthophora*) can profitably exploit patchy floral resources embedded in a shady habitat matrix such as, for instance, the sparse blooming individuals of rupicolous species hanging from rock cliffs. The high-*K*, irradiance-dependent ectothermic bees, in contrast, should be mostly constrained to open, sunlit habitats.

### Implications for pollination

Daily, seasonal and habitat-type correlates of the thermal biology of bee assemblages are expected to have direct and indirect implications for their pollination service and the reproductive success of plants. Only the two presumably most consequential of these effects will be discussed here: the indirect effect resulting from spatio-temporal changes in the diversity of bees available to plants, and the direct effect arising from the relationships between thermal biology and flower visitation rate.

Bee diversity can enhance plant reproductive success through complementarity effects (Herrera 2000, Fründ et al. 2013b, Martins et al. 2015), hence its daily, seasonal and among-habitat variation shaped by thermal biology is expected to impinge on plant reproductive success. For example, the thermally-mediated daily rhythm in composition of bee assemblages will provide a backdrop, or template, that condition not only the composition, but also the diversity of the bee pollinator assemblages of individual plant species. Such effects will be mediated by the substantial variation among plant species in timing and duration of their daily period of flower presentation to pollinators. Only about half of entomophilous species in my study region have functional flowers presented continuously throughout daytime (e.g., Apiaceae, Lamiaceae, Primulaceae), while the rest have flowers that open during early (e.g., Cistaceae) or late morning (e.g., subfamily Cichorioideae in Asteraceae) and close, wither or are shed around noon or shortly thereafter (C. M. Herrera, *unpublished data*). Other things being equal, bee-pollinated plants whose flowers are continuously open will be able to exploit a thermally broader assortment of bee taxa than those with open flowers restricted to narrower daytime intervals (see, e.g., Schlising 1970, Linsley 1978, Willmer and Corbet 1981, Herrera 1990, Fründ et al. 2011, for circumstantial support to this prediction). By the same token, taxonomic, phylogenetic and (presumably) functional diversity of bees available to plants are expected to be narrower in shady, closed habitats than in open, sunlit ones because of habitat-related variation in thermal biology profiles of bee assemblages, which would eventually lead to differences among habitats in bee pollinator service and plant reproductive success (Grab et al. 2019).

A direct effect of the diversity in bee thermal biology on plant reproduction is expected from the relationships between thermal biology parameters and flower visitation rates, as the latter is a key determinant of the “quantity” component in plant-pollinator relationships (Herrera 1989, Utelli and Roy 2000, Alonso et al. 2012, Rocca and Sazima 2013). After accounting for potentially confounding factors (body mass, flower abundance, plant species), flower visitation rate was found here to be inversely related to *K*, and directly related to *T*_th_ and *T*_exc_, but only among those plant species with restrictive flowers. This finding should be interpreted in the light of the direct relationship linking thoracic temperature with physiological performance and muscular power output in bees (Esch 1976, 1988, Casey 1988, Coelho 1991, Sinclair et al. 2016). When faced with flowers whose rewards are hidden and/or hard to reach, the greater muscular power and foraging speed of warmer bees enabled them to achieve higher visitation rates than cooler bees, but they had no advantage when visiting flowers with open, nonrestrictive perianths. Due to this interaction between perianth type and thermal parameters, open flowers were visited most often by bees belonging to the cool-bodied evolutionary lineage (high *K* and low *T*_th_ and *T*_exc_), while the reverse held for restrictive flowers (Figure 10). Through this mechanism, plants with contrasting floral morphologies could be partitioning bee pollinators on the basis of the latter’s thermal biology and, eventually, evolutionary origin.

## CONCLUDING REMARKS

Most research on bee thermal biology has historically focused on a few, phylogenetically closely-related lineages of large bees in the family Apidae possessing remarkable endothermic and thermoregulatory abilities (mostly species in the genera *Anthophora*, *Apis*, *Bombus* and *Xylocopa*) and, as a consequence, general statements on the thermal biology of bees have tended to emphasize the importance of these particular thermal biology features for bees and their interactions with plants (Herrera et al. 2023). By examining a large, taxonomically and phylogenetically diverse sample of bees encompassing virtually all bee pollinators of a large regional community of entomophilous plants, the present study has documented the true extent of thermal biology diversity involved in a natural plant-bee pollination system. Although infrequently undertaken in recent years, community-wide studies focusing on the pollination of large samples of plant species served first to set the foundations of modern pollination ecology (Müller 1883, Robertson 1895, 1924, 1929), and then to deepening our understanding of the ecology and evolution of plant-pollinator interactions (Arroyo et al. 1982, 1985, Inouye and Pyke 1988, McCall and Primack 1992, Lázaro et al. 2013, Herrera 2020). The present study went one step further in this tradition by adding another community-wide layer of information to the plant-pollinator system, namely the one involving the thermal biology of virtually the whole assemblage of bee pollinators, so that roughly the entire plant-bee pollination system of the region was eventually under scrutiny. Results not only confirm Herrera et al.’s (2023) contention that the thermal biology diversity of bee communities should be much broader than sometimes inferred from thermal biology studies based on taxonomically restricted bee samples, but also that bee thermal diversity can lie at the very core of the evolution and broad spatio-temporal organization of the entire multispecies plant-bee pollinator assemblage.

## ACKNOWLEDGMENTS

I’m deeply indebted to Oscar Aguado, Enrique Asensio, Leopoldo Castro, Curro Molina, Andreas Müller, Alejandro Núñez, Concepción Ornosa, Javier Ortiz, Max Kasparek, Klaus Schönitzer, Jan Smit and Thomas Wood for identification of bee specimens. Consejería de Medio Ambiente, Junta de Andalucía, granted permission to work in the Sierra de Cazorla and provided invaluable facilities there. Thanks are also due to Mónica Medrano for encouragement and discussions over so many years of pollinator censusing, and also for suggestions on the manuscript. The research reported in this paper received no specific grant from any funding agency.

## DATA AVAILABILITY STATEMENT

Data, metadata and R script for the analyses have been deposited at figshare (https://figshare.com/s/adef8af1e1dd57a2a665)

