## Appendix S1 for "Thermal biology diversity of bee pollinators: taxonomic, phylogenetic and plant community-level correlates"

TABLE S1 List of the 95 species of bees with paired field measurements of thoracic temperature ($T_{th}$) and air temperature at the spot of capture ($T_{air}$)

| | | $T_{th}$ (ºC) | | | $T_{air}$ (ºC) | |
|---|---|---|---|---|---|---|
| Species | *N* | Mean | SE | | Mean | SE |
| *Amegilla albigena* | 27 | 39.27 | 0.32 | | 31.00 | 0.71 |
| *Amegilla ochroleuca* | 22 | 39.48 | 0.55 | | 26.34 | 0.83 |
| *Amegilla quadrifasciata* | 16 | 41.19 | 0.52 | | 29.47 | 0.92 |
| *Andrena antigana* | 24 | 27.96 | 0.42 | | 19.93 | 0.41 |
| *Andrena assimilis* | 23 | 27.96 | 0.46 | | 13.08 | 0.50 |
| *Andrena baetica* | 39 | 28.78 | 0.26 | | 21.82 | 0.32 |
| *Andrena bicolor* | 71 | 26.68 | 0.24 | | 19.49 | 0.39 |
| *Andrena bimaculata* | 2 | 26.00 | 0.60 | | 15.50 | 0.10 |
| *Andrena chrysosceles* | 1 | 27.50 | | | 18.20 | |
| *Andrena cyanomicans* | 1 | 28.30 | | | 20.10 | |
| *Andrena flavipes* | 43 | 28.92 | 0.44 | | 17.80 | 0.57 |
| *Andrena fulva* | 24 | 27.59 | 0.70 | | 16.71 | 0.97 |
| *Andrena fulvicornis* | 1 | 30.10 | | | 23.30 | |
| *Andrena humilis* | 2 | 24.10 | 1.30 | | 17.15 | 1.45 |
| *Andrena impressa* | 1 | 24.30 | | | 14.40 | |
| *Andrena labialis* | 32 | 31.26 | 0.36 | | 21.22 | 0.37 |
| *Andrena lagopus* | 40 | 29.53 | 0.39 | | 20.26 | 0.28 |
| *Andrena leptopyga* | 1 | 31.40 | | | 22.50 | |
| *Andrena livens* | 27 | 25.59 | 0.44 | | 17.89 | 0.35 |
| *Andrena longibarbis* | 1 | 31.40 | | | 23.00 | |
| *Andrena nigroaenea* | 28 | 28.17 | 0.62 | | 17.86 | 0.64 |

| | | | | | | |
|---|---|---|---|---|---|---|
| *Andrena pilipes* | 3 | 31.33 | 0.69 | | 21.57 | 1.40 |
| *Andrena rhenana* | 15 | 29.22 | 0.51 | | 22.13 | 0.44 |
| *Andrena rhyssonota* | 42 | 26.99 | 0.34 | | 19.17 | 0.38 |
| *Andrena sardoa* | 29 | 28.80 | 0.58 | | 21.08 | 0.42 |
| *Andrena schencki* | 1 | 33.00 | | | 26.00 | |
| *Andrena suerinensis* | 1 | 28.30 | | | 19.70 | |
| *Andrena thoracica* | 1 | 28.50 | | | 15.30 | |
| *Andrena tibialis* | 3 | 23.10 | 0.90 | | 15.13 | 0.74 |
| *Andrena trimmerana* | 39 | 28.81 | 0.45 | | 18.77 | 0.64 |
| *Andrena vetula* | 1 | 27.90 | | | 20.20 | |
| *Andrena villipes* | 29 | 26.01 | 0.42 | | 15.50 | 0.27 |
| *Andrena vulpecula* | 1 | 34.00 | | | 27.10 | |
| *Anthidiellum breviusculum* | 39 | 34.61 | 0.52 | | 31.99 | 0.52 |
| *Anthidium florentinum* | 81 | 38.86 | 0.26 | | 29.57 | 0.45 |
| *Anthidium manicatum* | 1 | 39.90 | | | 28.30 | |
| *Anthophora aestivalis* | 3 | 38.60 | 2.36 | | 19.40 | 1.99 |
| *Anthophora affinis* | 1 | 39.20 | | | 19.80 | |
| *Anthophora atroalba* | 3 | 37.00 | 2.41 | | 20.13 | 1.59 |
| *Anthophora canescens* | 1 | 38.60 | | | 18.50 | |
| *Anthophora crassipes* | 35 | 38.45 | 0.28 | | 27.08 | 0.56 |
| *Anthophora dispar* | 85 | 38.65 | 0.32 | | 16.61 | 0.46 |
| *Anthophora fulvodimidiata* | 1 | 34.30 | | | 21.90 | |
| *Anthophora leucophaea* | 3 | 39.50 | 1.17 | | 17.40 | 2.17 |
| *Anthophora plumipes* | 11 | 35.05 | 0.56 | | 15.73 | 0.94 |
| *Anthophora pubescens* | 1 | 36.00 | | | 26.60 | |
| *Anthophora retusa* | 9 | 37.84 | 0.83 | | 19.48 | 0.96 |
| *Anthophora romandii* | 14 | 39.18 | 0.56 | | 18.77 | 0.81 |
| *Apis mellifera* | 243 | 38.76 | 0.23 | | 25.27 | 0.49 |
| *Bombus pascuorum* | 58 | 37.17 | 0.46 | | 24.14 | 0.67 |
| *Bombus pratorum* | 34 | 36.89 | 0.36 | | 14.06 | 0.83 |

| | | | | | | |
|---|---|---|---|---|---|---|
| *Bombus terrestris* | 152 | 38.62 | 0.25 | | 24.47 | 0.52 |
| *Bombus vestalis* | 3 | 36.57 | 0.52 | | 16.57 | 1.90 |
| *Ceratina cucurbitina* | 1 | 28.70 | | | 27.90 | |
| *Ceratina mocsaryi* | 18 | 34.83 | 0.78 | | 32.09 | 0.77 |
| *Colletes acutiformis* | 23 | 29.86 | 0.49 | | 19.32 | 0.40 |
| *Colletes acutus* | 4 | 30.90 | 1.29 | | 21.25 | 1.84 |
| *Colletes albomaculatus* | 1 | 24.20 | | | 17.20 | |
| *Colletes cunicularius* | 29 | 30.44 | 0.45 | | 18.81 | 0.63 |
| *Colletes hederae* | 100 | 29.85 | 0.30 | | 23.46 | 0.33 |
| *Colletes nigricans* | 1 | 29.30 | | | 26.00 | |
| *Colletes sierrensis* | 2 | 27.10 | 1.10 | | 23.40 | 0.10 |
| *Eucera chrysopyga* | 3 | 30.07 | 3.06 | | 18.87 | 0.98 |
| *Eucera collaris* | 21 | 31.73 | 0.59 | | 19.56 | 0.44 |
| *Eucera decolorata* | 10 | 29.46 | 0.63 | | 18.17 | 0.68 |
| *Eucera elongatula* | 26 | 29.55 | 0.45 | | 19.27 | 0.19 |
| *Eucera eucnemidea* | 2 | 27.75 | 1.15 | | 16.85 | 0.55 |
| *Eucera hispaliensis* | 13 | 31.72 | 0.48 | | 18.92 | 0.43 |
| *Eucera nigrilabris* | 5 | 34.56 | 1.85 | | 20.08 | 1.52 |
| *Halictus fulvipes* | 1 | 23.00 | | | 20.30 | |
| *Halictus quadricinctus* | 1 | 28.00 | | | 17.20 | |
| *Halictus scabiosae* | 13 | 29.51 | 0.72 | | 23.52 | 1.15 |
| *Icteranthidium laterale* | 2 | 31.80 | 0.60 | | 25.55 | 0.05 |
| *Lasioglossum immunitum* | 26 | 24.90 | 0.41 | | 18.32 | 0.37 |
| *Lasioglossum leucozonium* | 1 | 21.50 | | | 17.00 | |
| *Lasioglossum malachurum* | 4 | 24.85 | 1.08 | | 17.98 | 0.27 |
| *Lasioglossum perclavipes* | 1 | 25.80 | | | 19.60 | |
| *Lasioglossum xanthopus* | 1 | 24.10 | | | 18.40 | |
| *Lithurgus chrysurus* | 60 | 35.73 | 0.32 | | 27.77 | 0.46 |
| *Megachile albisecta* | 43 | 35.98 | 0.41 | | 27.60 | 0.58 |
| *Megachile centuncularis* | 4 | 32.05 | 1.60 | | 23.78 | 2.51 |

| | | | | | | |
|---|---|---|---|---|---|---|
| *Megachile octosignata* | 4 | 29.98 | 1.03 | | 21.48 | 1.18 |
| *Megachile pilicrus* | 2 | 32.15 | 0.85 | | 24.10 | 1.50 |
| *Megachile pilidens* | 23 | 34.26 | 0.82 | | 28.42 | 0.69 |
| *Megachile sicula* | 5 | 35.68 | 0.95 | | 20.24 | 0.94 |
| *Melecta albifrons* | 2 | 31.85 | 0.55 | | 17.35 | 2.15 |
| *Nomada succincta* | 1 | 28.50 | | | 12.90 | |
| *Osmia aurulenta* | 1 | 32.60 | | | 24.60 | |
| *Osmia bicornis* | 9 | 36.66 | 0.68 | | 20.18 | 1.09 |
| *Osmia cornuta* | 17 | 32.84 | 0.70 | | 15.02 | 0.73 |
| *Panurgus perezi* | 3 | 25.57 | 1.25 | | 16.87 | 0.58 |
| *Rhodanthidium sticticum* | 3 | 35.87 | 0.24 | | 22.27 | 0.27 |
| *Xylocopa cantabrita* | 35 | 39.19 | 0.36 | | 19.48 | 0.61 |
| *Xylocopa valga* | 5 | 41.88 | 1.04 | | 21.66 | 1.94 |
| *Xylocopa violacea* | 41 | 40.46 | 0.53 | | 25.10 | 0.92 |

TABLE S2 List of the 169 species of bees for which the warming constant *K* was experimentally estimated in the laboratory, number of individuals assayed (*N*), species means for *K*, and mean body mass of the individuals used in the experiments.

| | | *K* ($s^{-1}$) | | | Body mass (mg) | |
|---|---|---|---|---|---|---|
| Bee species | *N* | Mean | SE | | Mean | SE |
| *Amegilla albigena* | 10 | 0.01422 | 0.00151 | | 62.4 | 8.1 |
| *Amegilla quadrifasciata* | 5 | 0.00902 | 0.00101 | | 117.0 | 28.0 |
| *Andrena afzeliella* | 1 | 0.03277 | | | 17.8 | |
| *Andrena albopunctata* | 1 | 0.00834 | | | 137.0 | |
| *Andrena apicata* | 1 | 0.03347 | | | 16.5 | |
| *Andrena asperrima* | 3 | 0.01675 | 0.00290 | | 53.0 | 15.0 |
| *Andrena assimilis* | 4 | 0.01026 | 0.00134 | | 106.3 | 23.5 |
| *Andrena baetica* | 4 | 0.02113 | 0.00104 | | 28.8 | 1.4 |
| *Andrena bicolor* | 16 | 0.02797 | 0.00276 | | 26.2 | 2.2 |
| *Andrena bimaculata* | 1 | 0.02210 | | | 33.7 | |
| *Andrena cinerea* | 1 | 0.03221 | | | 21.0 | |
| *Andrena congruens* | 2 | 0.01564 | 0.00149 | | 52.7 | 5.4 |
| *Andrena djelfensis* | 5 | 0.06506 | 0.00240 | | 6.7 | 0.3 |
| *Andrena flavipes* | 14 | 0.02388 | 0.00214 | | 37.1 | 6.9 |
| *Andrena florentina* | 4 | 0.02271 | 0.00334 | | 32.4 | 8.9 |
| *Andrena fulva* | 3 | 0.01220 | 0.00052 | | 81.4 | 2.8 |
| *Andrena haemorrhoa* | 9 | 0.02424 | 0.00241 | | 31.9 | 5.0 |
| *Andrena hattorfiana* | 2 | 0.00994 | 0.00019 | | 117.5 | 5.4 |
| *Andrena helvola* | 2 | 0.04264 | 0.00449 | | 15.7 | 1.2 |
| *Andrena humilis* | 5 | 0.02192 | 0.00131 | | 29.4 | 2.9 |
| *Andrena impressa* | 16 | 0.02735 | 0.00210 | | 24.3 | 2.4 |
| *Andrena labialis* | 6 | 0.01811 | 0.00168 | | 45.5 | 6.5 |
| *Andrena lecana* | 1 | 0.04206 | | | 13.0 | |
| *Andrena lepida* | 4 | 0.02273 | 0.00450 | | 33.8 | 6.3 |

| | | | | | | |
|---|---|---|---|---|---|---|
| *Andrena leucophaea* | 3 | 0.03026 | 0.00041 | | 20.4 | 0.6 |
| *Andrena limata* | 1 | 0.01139 | | | 94.3 | |
| *Andrena livens* | 7 | 0.01813 | 0.00073 | | 36.6 | 1.3 |
| *Andrena medeninensis* | 2 | 0.02318 | 0.00072 | | 26.6 | 1.2 |
| *Andrena melacana* | 1 | 0.03047 | | | 20.5 | |
| *Andrena minutula* | 5 | 0.05111 | 0.00484 | | 8.0 | 1.2 |
| *Andrena morio* | 3 | 0.00955 | 0.00030 | | 133.6 | 6.0 |
| *Andrena nana* | 1 | 0.05941 | | | 6.7 | |
| *Andrena nigroaenea* | 21 | 0.01728 | 0.00122 | | 56.2 | 6.6 |
| *Andrena orbitalis* | 1 | 0.02438 | | | 26.2 | |
| *Andrena ovatula* | 3 | 0.01806 | 0.00110 | | 43.1 | 1.5 |
| *Andrena pandellei* | 1 | 0.02371 | | | 32.1 | |
| *Andrena pilipes* | 6 | 0.01340 | 0.00143 | | 79.2 | 15.4 |
| *Andrena propinqua* | 2 | 0.01801 | 0.00118 | | 47.4 | 0.3 |
| *Andrena ranunculi* | 1 | 0.02778 | | | 27.8 | |
| *Andrena rhyssonota* | 2 | 0.02116 | 0.00485 | | 35.5 | 10.9 |
| *Andrena sardoa* | 4 | 0.02259 | 0.00559 | | 40.3 | 8.7 |
| *Andrena schencki* | 1 | 0.01449 | | | 73.7 | |
| *Andrena senecionis* | 1 | 0.02688 | | | 23.9 | |
| *Andrena synadelpha* | 2 | 0.01943 | 0.00280 | | 42.5 | 8.5 |
| *Andrena thoracica* | 3 | 0.00992 | 0.00012 | | 116.4 | 8.3 |
| *Andrena tibialis* | 28 | 0.01796 | 0.00074 | | 48.9 | 3.6 |
| *Andrena trimmerana* | 21 | 0.01880 | 0.00169 | | 54.4 | 6.3 |
| *Andrena villipes* | 3 | 0.01811 | 0.00360 | | 41.3 | 13.3 |
| *Anthidiellum breviusculum* | 3 | 0.02778 | 0.00197 | | 20.2 | 2.9 |
| *Anthidiellum strigatum* | 9 | 0.02876 | 0.00051 | | 19.6 | 1.0 |
| *Anthidium cingulatum* | 1 | 0.01054 | | | 84.2 | |
| *Anthidium diadema* | 1 | 0.00825 | | | 106.4 | |
| *Anthidium florentinum* | 4 | 0.00704 | 0.00048 | | 153.9 | 20.0 |
| *Anthidium manicatum* | 5 | 0.00791 | 0.00088 | | 127.8 | 20.8 |

| | | | | | | |
|---|---|---|---|---|---|---|
| *Anthidium taeniatum* | 1 | 0.01685 | | | 42.6 | |
| *Anthophora aestivalis* | 1 | 0.00879 | | | 90.1 | |
| *Anthophora atroalba* | 1 | 0.00663 | | | 162.7 | |
| *Anthophora bimaculata* | 1 | 0.02117 | | | 28.1 | |
| *Anthophora canescens* | 11 | 0.00659 | 0.00052 | | 185.5 | 24.5 |
| *Anthophora crassipes* | 7 | 0.01022 | 0.00111 | | 88.1 | 7.4 |
| *Anthophora dispar* | 6 | 0.00738 | 0.00018 | | 145.9 | 9.1 |
| *Anthophora femorata* | 4 | 0.00617 | 0.00051 | | 182.7 | 11.7 |
| *Anthophora fulvitarsis* | 1 | 0.00551 | | | 199.8 | |
| *Anthophora fulvodimidiata* | 7 | 0.01078 | 0.00049 | | 74.5 | 3.5 |
| *Anthophora leucophaea* | 11 | 0.00947 | 0.00038 | | 92.1 | 8.3 |
| *Anthophora plumipes* | 2 | 0.00747 | 0.00098 | | 135.8 | 37.1 |
| *Anthophora pubescens* | 1 | 0.01424 | | | 51.8 | |
| *Anthophora retusa* | 10 | 0.00800 | 0.00024 | | 111.0 | 4.2 |
| *Anthophora romandii* | 1 | 0.00944 | | | 85.7 | |
| *Apis mellifera* | 3 | 0.01090 | 0.00019 | | 80.6 | 3.0 |
| *Bombus pascuorum* | 3 | 0.00719 | 0.00035 | | 147.8 | 13.2 |
| *Bombus pratorum* | 8 | 0.00664 | 0.00034 | | 154.8 | 10.6 |
| *Bombus terrestris* | 7 | 0.00705 | 0.00094 | | 234.1 | 64.3 |
| *Bombus vestalis* | 2 | 0.00691 | 0.00001 | | 207.1 | 15.7 |
| *Ceratina chalybea* | 2 | 0.03307 | 0.00196 | | 17.2 | 0.2 |
| *Ceratina cucurbitina* | 1 | 0.06801 | | | 5.1 | |
| *Ceratina cyanea* | 6 | 0.05194 | 0.00234 | | 9.3 | 0.7 |
| *Ceratina gravidula* | 4 | 0.03114 | 0.00250 | | 23.7 | 3.0 |
| *Ceratina mocsaryi* | 2 | 0.02983 | 0.00335 | | 19.8 | 2.9 |
| *Chelostoma emarginatum* | 4 | 0.03168 | 0.00318 | | 19.1 | 2.1 |
| *Chelostoma florisomne* | 4 | 0.03395 | 0.00204 | | 18.0 | 2.2 |
| *Coelioxys acanthura* | 1 | 0.01650 | | | 42.8 | |
| *Coelioxys brevis* | 1 | 0.02806 | | | 19.7 | |
| *Colletes abeillei* | 2 | 0.01330 | 0.00070 | | 64.1 | 4.5 |

| | | | | | | |
|---|---|---|---|---|---|---|
| *Colletes cunicularius* | 11 | 0.01260 | 0.00096 | | 75.3 | 8.8 |
| *Colletes hederae* | 9 | 0.01805 | 0.00172 | | 44.7 | 4.8 |
| *Colletes nigricans* | 3 | 0.01649 | 0.00024 | | 45.0 | 3.1 |
| *Colletes sierrensis* | 8 | 0.02750 | 0.00096 | | 22.0 | 1.2 |
| *Epeolus fallax* | 3 | 0.03056 | 0.00307 | | 18.7 | 2.6 |
| *Eucera caspica* | 4 | 0.01123 | 0.00065 | | 78.0 | 11.5 |
| *Eucera clypeata* | 1 | 0.01549 | | | 42.1 | |
| *Eucera elongatula* | 12 | 0.01700 | 0.00046 | | 37.5 | 2.1 |
| *Eucera eucnemidea* | 9 | 0.01468 | 0.00042 | | 57.5 | 3.1 |
| *Eucera nigrifacies* | 2 | 0.01772 | 0.00130 | | 46.8 | 2.3 |
| *Eucera nigrilabris* | 3 | 0.00925 | 0.00075 | | 106.3 | 11.1 |
| *Halictus fulvipes* | 4 | 0.01863 | 0.00099 | | 42.9 | 6.5 |
| *Halictus gemmeus* | 6 | 0.05358 | 0.00373 | | 6.6 | 0.3 |
| *Halictus pollinosus* | 1 | 0.03603 | | | 22.0 | |
| *Halictus quadricinctus* | 3 | 0.00940 | 0.00153 | | 119.6 | 18.7 |
| *Halictus scabiosae* | 2 | 0.01527 | 0.00158 | | 64.3 | 17.1 |
| *Halictus smaragdulus* | 5 | 0.04329 | 0.00317 | | 7.5 | 1.0 |
| *Halictus subauratus* | 3 | 0.03733 | 0.00836 | | 18.3 | 5.2 |
| *Halictus tridivisus* | 1 | 0.02114 | | | 33.4 | |
| *Heriades crenulatus* | 4 | 0.03761 | 0.00345 | | 13.2 | 1.2 |
| *Hoplitis benoisti* | 2 | 0.02079 | 0.00137 | | 30.9 | 2.3 |
| *Hoplitis curvipes* | 1 | 0.01144 | | | 80.2 | |
| *Hylaeus communis* | 4 | 0.06233 | 0.00524 | | 5.6 | 0.5 |
| *Hylaeus gibbus* | 1 | 0.04232 | | | 8.5 | |
| *Hylaeus variegatus* | 7 | 0.04768 | 0.00545 | | 10.6 | 1.3 |
| *Icteranthidium grohmanni* | 3 | 0.01785 | 0.00321 | | 45.7 | 14.5 |
| *Icteranthidium laterale* | 3 | 0.00838 | 0.00071 | | 128.8 | 16.0 |
| *Lasioglossum bimaculatum* | 1 | 0.03286 | | | 19.4 | |
| *Lasioglossum brevicorne* | 1 | 0.04616 | | | 8.4 | |
| *Lasioglossum buccale* | 1 | 0.03469 | | | 18.8 | |

| | | | | | | |
|---|---|---|---|---|---|---|
| *Lasioglossum calceatum* | 1 | 0.02505 | | | 26.4 | |
| *Lasioglossum laevigatum* | 2 | 0.02689 | 0.00299 | | 26.4 | 1.8 |
| *Lasioglossum marginatum* | 3 | 0.03343 | 0.00231 | | 16.5 | 1.7 |
| *Lasioglossum mediterraneum* | 9 | 0.03609 | 0.00140 | | 16.0 | 1.0 |
| *Lasioglossum minutissimum* | 1 | 0.04882 | | | 9.2 | |
| *Lasioglossum pallens* | 5 | 0.05246 | 0.00296 | | 9.1 | 0.3 |
| *Lasioglossum pauperatum* | 1 | 0.07176 | | | 6.3 | |
| *Lasioglossum punctatissimum* | 2 | 0.05425 | 0.00164 | | 8.6 | 0.3 |
| *Lasioglossum puncticolle* | 1 | 0.04741 | | | 11.6 | |
| *Lasioglossum quadrinotatum* | 1 | 0.05159 | | | 9.8 | |
| *Lasioglossum sphecodimorphum* | 2 | 0.06403 | 0.00852 | | 7.2 | 1.7 |
| *Lasioglossum subhirtum* | 1 | 0.03130 | | | 17.7 | |
| *Lasioglossum transitorium* | 1 | 0.05187 | | | 12.4 | |
| *Lasioglossum villosulum* | 1 | 0.04718 | | | 10.3 | |
| *Lasioglossum xanthopus* | 5 | 0.01811 | 0.00224 | | 51.2 | 5.8 |
| *Lithurgus chrysurus* | 3 | 0.01152 | 0.00104 | | 76.6 | 6.9 |
| *Megachile albisecta* | 5 | 0.00947 | 0.00110 | | 105.6 | 15.1 |
| *Megachile albonotata* | 2 | 0.01061 | 0.00242 | | 94.5 | 27.9 |
| *Megachile apicalis* | 2 | 0.01911 | 0.00061 | | 33.6 | 0.8 |
| *Megachile centuncularis* | 4 | 0.01495 | 0.00139 | | 52.3 | 7.7 |
| *Megachile giraudi* | 1 | 0.01464 | | | 52.0 | |
| *Megachile lagopoda* | 2 | 0.00601 | 0.00006 | | 177.9 | 16.4 |
| *Megachile octosignata* | 1 | 0.01973 | | | 28.8 | |
| *Megachile pilidens* | 11 | 0.01940 | 0.00096 | | 33.7 | 2.6 |
| *Megachile pyrenaica* | 2 | 0.00850 | 0.00062 | | 119.3 | 22.0 |
| *Megachile rotundata* | 1 | 0.01785 | | | 43.8 | |
| *Melecta albifrons* | 2 | 0.01101 | 0.00126 | | 78.1 | 10.7 |
| *Melecta italica* | 2 | 0.01028 | 0.00049 | | 91.4 | 2.1 |
| *Melecta luctuosa* | 3 | 0.00851 | 0.00030 | | 118.1 | 9.7 |
| *Nomada bifasciata* | 6 | 0.02487 | 0.00209 | | 33.6 | 3.9 |

| | | | | | | |
|---|---|---|---|---|---|---|
| *Nomada flavilabris* | 1 | 0.03186 | | | 17.4 | |
| *Nomada flavoguttata* | 2 | 0.03900 | 0.00790 | | 12.8 | 3.3 |
| *Nomada hispanica* | 1 | 0.03388 | | | 17.3 | |
| *Nomada marshamella* | 1 | 0.02036 | | | 36.5 | |
| *Nomada melathoracica* | 4 | 0.02176 | 0.00301 | | 42.8 | 11.4 |
| *Nomada ruficornis* | 1 | 0.03605 | | | 18.6 | |
| *Nomada succincta* | 5 | 0.01881 | 0.00149 | | 42.0 | 3.3 |
| *Osmia argyropyga* | 1 | 0.01948 | | | 36.0 | |
| *Osmia aurulenta* | 1 | 0.01111 | | | 73.7 | |
| *Osmia bicornis* | 4 | 0.01497 | 0.00143 | | 51.3 | 8.8 |
| *Osmia cornuta* | 5 | 0.01055 | 0.00138 | | 89.8 | 15.2 |
| *Osmia niveocincta* | 1 | 0.01727 | | | 49.4 | |
| *Osmia submicans* | 2 | 0.02403 | 0.00033 | | 25.4 | 2.4 |
| *Osmia tricornis* | 1 | 0.01018 | | | 89.6 | |
| *Panurgus banksianus* | 7 | 0.02417 | 0.00121 | | 25.5 | 1.6 |
| *Panurgus calcaratus* | 5 | 0.04543 | 0.00194 | | 9.0 | 0.3 |
| *Protosmia capitata* | 4 | 0.04366 | 0.00539 | | 15.8 | 3.1 |
| *Rhodanthidium sticticum* | 2 | 0.00865 | 0.00024 | | 124.4 | 0.4 |
| *Sphecodes albilabris* | 3 | 0.01617 | 0.00311 | | 59.2 | 13.7 |
| *Sphecodes monilicornis* | 1 | 0.02035 | | | 34.6 | |
| *Thyreus ramosus* | 2 | 0.01684 | 0.00068 | | 43.1 | 4.1 |
| *Thyreus truncatus* | 1 | 0.01155 | | | 86.6 | |
| *Xylocopa cantabrita* | 2 | 0.00508 | 0.00034 | | 246.8 | 24.3 |
| *Xylocopa iris* | 1 | 0.00784 | | | 167.1 | |
| *Xylocopa violacea* | 5 | 0.00437 | 0.00014 | | 462.7 | 50.9 |

TABLE S3 Results of partitioning the variance among the individual bees sampled in warming constant (*K*), thoracic temperature ($T_{th}$) and thermal excess ($T_{exc}$), using a hierarchically nested model with genus nested within family, species nested within genus, and residual variance accounting for variance among individuals of the same species. To facilitate comparisons figures are percentages with respect to the total variance. *N* = number of individual measurements.

| Source of variance | *K* (*N* = 645) | $T_{th}$ (*N* = 1936) | $T_{exc}$ (*N* = 1936) |
|---|---|---|---|
| Family | 9.73 | 5.08 | 14.21 |
| Genus | 42.97 | 22.96 | 40.95 |
| Species | 35.60 | 50.30 | 17.58 |
| Residual (among individuals within species) | 11.70 | 21.66 | 27.26 |

TABLE S4 Summary of results of generalized aditive models testing for the effects of Julian date (days from 1 January; nonparametric smooth term), body mass (mg, log-transformed), bee family, and bee genus (nested within family) on the warming constant (*K*, log-transformed), thoracic temperature ($T_{th}$, ºC) and thermal excess ($T_{exc} = T_{th} – T_{air}$, ºC) of the bee assemblages recorded in pollinator censuses.

| Response variable | Predictor | Degrees of freedom | *F* | *P*-value |
|---|---|---|---|---|
| $Log_{10}$ (*K*) | smooth(Julian date) | 8.7, 8667.3 | 93.1 | <2.2E-16 |
| | $log_{10}$ (body mass) | 1, 8667.3 | 4951 | <2.2E-16 |
| | Bee family | 4, 8667.3 | 11378 | <2.2E-16 |
| | Bee genus | 18, 8667.3 | 9318 | <2.2E-16 |
| $T_{th}$ | smooth(Julian date) | 22.9, 8928.0 | 99.2 | <2.2E-16 |
| | $log_{10}$ (body mass) | 1, 8928.0 | 3857 | <2.2E-16 |
| | Bee family | 4, 8928.0 | 5770 | <2.2E-16 |
| | Bee genus | 19, 8928.0 | 8241 | <2.2E-16 |
| $T_{exc}$ | smooth(Julian date) | 22.9, 8928.0 | 214.1 | <2.2E-16 |
| | $log_{10}$ (body mass) | 1, 8928.0 | 16.6 | 4.7E-05 |
| | Bee family | 4, 8928.0 | 148.3 | <2.2E-16 |
| | Bee genus | 19, 8928.0 | 649.2 | <2.2E-16 |

TABLE S5 Number of bee individuals and species recorded in censuses to which thermal biology data (species' means for $K$, $T_{th}$ and $T_{exc}$) could be assigned, shown separately for habitat types.

| | Bee individuals with assigned thermal biology data | | | Bee species with assigned thermal biology data | |
|---|---|---|---|---|---|
| Habitat type | $K$ | $T_{th}$, $T_{exc}$ | | $K$ | $T_{th}$, $T_{exc}$ |
| Disturbances | 2845 | 1531 | | 74 | 43 |
| Dolomitic outcrops | 767 | 295 | | 53 | 28 |
| Dwarf mountain scrub | 2293 | 1003 | | 87 | 44 |
| Forest clearings and edges | 2236 | 1378 | | 75 | 44 |
| Forest interior | 964 | 648 | | 41 | 28 |
| Grasslands and meadows | 1824 | 1157 | | 74 | 44 |
| Rock cliffs | 357 | 298 | | 25 | 14 |
| Streams/springs | 1392 | 844 | | 63 | 40 |
| Tall sclerophyllous scrub | 2243 | 1858 | | 60 | 41 |
| Total | 14921 | 9012 | | 134 | 71 |

TABLE S6 Summary of results of linear models testing for the effects of habitat type, body mass (mg, log-transformed), bee family, and bee genus (nested within family) on the warming constant (*K*, log-transformed), thoracic temperature ($T_{th}$, ºC) and thermal excess ($T_{exc} = T_{th} – T_{air}$, ºC) of bee assemblages recorded in pollinator censuses, all plant species combined within each habitat type. *N* = 8,696 (*K* analysis) and 8,951 ($T_{th}$ and $T_{exc}$ analyses) complete individual data.

| Response variable | Predictor | Degrees of freedom | *F* | *P*-value |
|---|---|---|---|---|
| $Log_{10}$ (*K*) | Habitat type | 8, 8668 | 36.7 | <2.2E-16 |
| | $log_{10}$ (body mass) | 1, 8668 | 4614.6 | <2.2E-16 |
| | Bee family | 4, 8668 | 743.3 | <2.2E-16 |
| | Bee genus | 14, 8668 | 1203.5 | <2.2E-16 |
| $T_{th}$ | Habitat type | 8, 8922 | 54.2 | <2.2E-16 |
| | $log_{10}$ (body mass) | 1, 8922 | 3979.5 | <2.2E-16 |
| | Bee family | 4, 8922 | 14124.9 | <2.2E-16 |
| | Bee genus | 15, 8922 | 1293.9 | <2.2E-16 |
| $T_{exc}$ | Habitat type | 8, 8922 | 51.1 | <2.2E-16 |
| | $log_{10}$ (body mass) | 1, 8922 | 22.0 | <2.8E-06 |
| | Bee family | 4, 8922 | 1088.7 | <2.2E-16 |
| | Bee genus | 15, 8922 | 614.6 | <2.2E-16 |

FIGURE S1. Relationship between warming constant *K* and body mass across the sample of 169 bee species studied (Table S2). Each symbol corresponds to a species, and the blue line is the least-squares fitted linear regression (shaded area is the 95% confidence interval). Note the logarithmic scales on both axes.

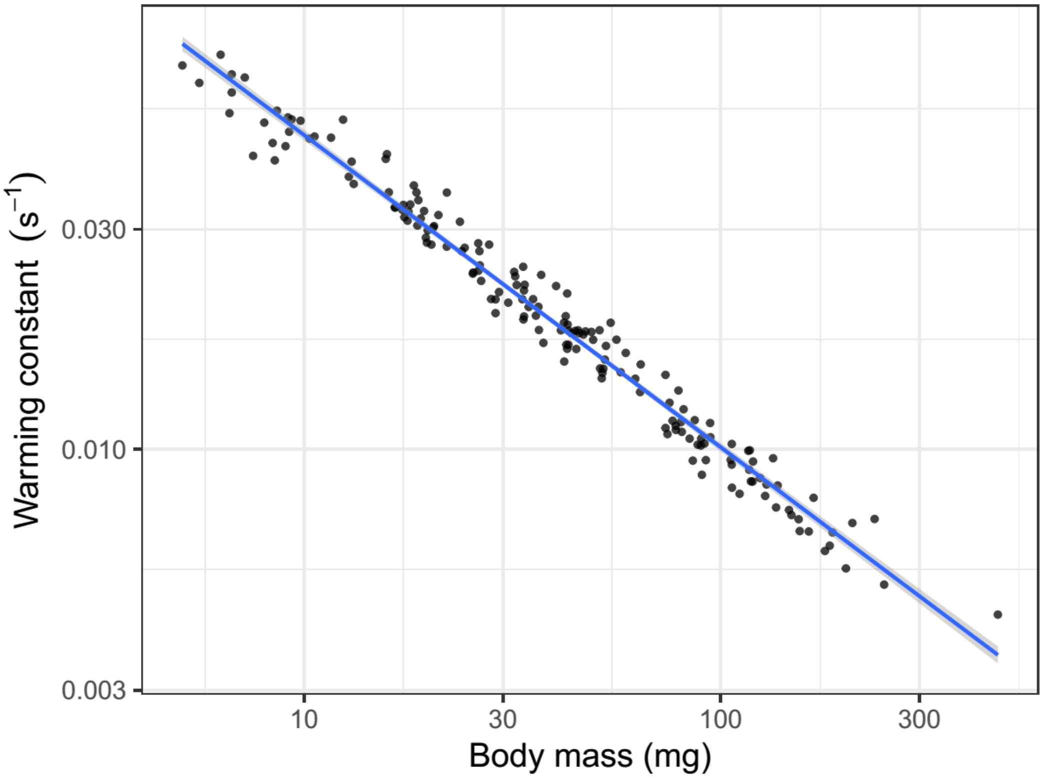

FIGURE S2. Relationship between warming constant *K* and body mass for the five bee families represented in the sample. Lines are least-squares fitted linear regressions. Note the logarithmic scales on both axes.

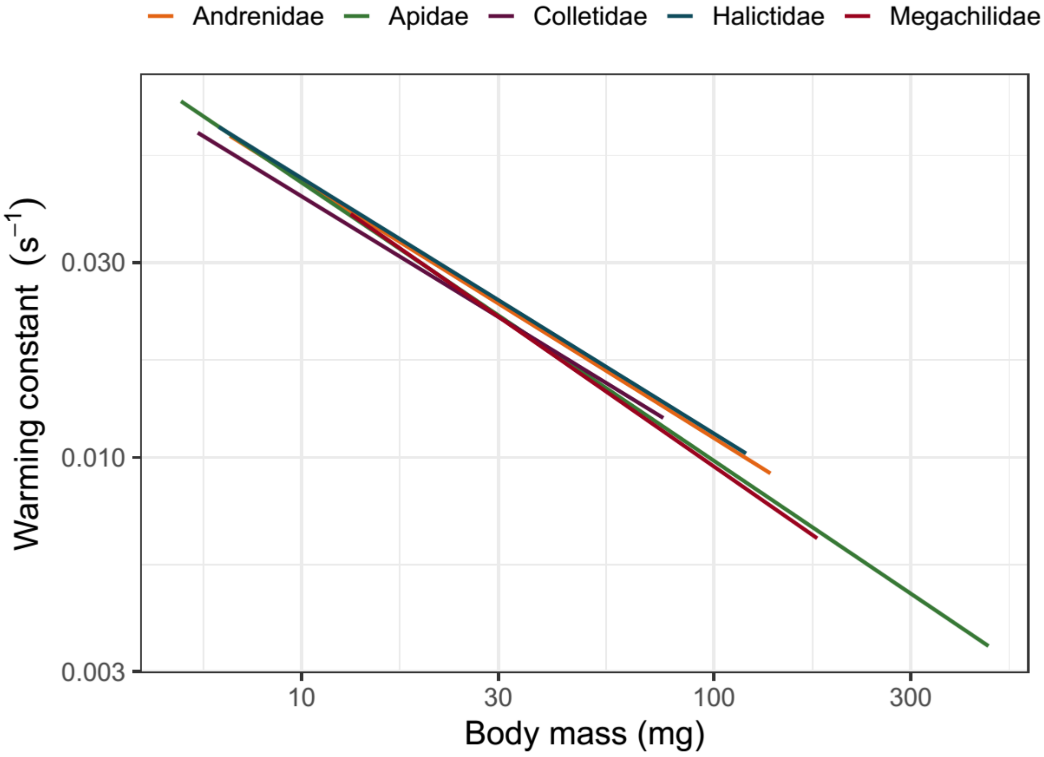
